# Symmetry, gauge freedoms, and the interpretability of sequence-function Relationships

**DOI:** 10.1101/2024.05.12.593774

**Authors:** Anna Posfai, David M. McCandlish, Justin B. Kinney

## Abstract

Quantitative models that describe how biological sequences encode functional activities are ubiquitous in modern biology. One important aspect of these models is that they commonly exhibit gauge freedoms, i.e., directions in parameter space that do not affect model predictions. In physics, gauge freedoms arise when physical theories are formulated in ways that respect fundamental symmetries. However, the connections that gauge freedoms in models of sequence-function relationships have to the symmetries of sequence space have yet to be systematically studied. In this work we study the gauge freedoms of models that respect a specific symmetry of sequence space: the group of position-specific character permutations. We find that gauge freedoms arise when model parameters transform under redundant irreducible matrix representations of this group. Based on this finding, we describe an “embedding distillation” procedure that enables both analytic calculation of the number of independent gauge freedoms and efficient computation of a sparse basis for the space of gauge freedoms. We also study how parameter transformation behavior affects parameter interpretability. We find that in many (and possibly all) nontrivial models, the ability to interpret individual model parameters as quantifying intrinsic allelic effects requires that gauge freedoms be present. This finding establishes an incompatibility between two distinct notions of parameter interpretability. Our work thus advances the understanding of symmetries, gauge freedoms, and parameter interpretability in models of sequence-function relationships.

## INTRODUCTION

Understanding the quantitative nature of sequence-function relationships is a major goal of modern biology [1]. To study a sequence-function relationship of interest, researchers often propose a mathematical model, fit the parameters of the model to data, then biologically interpret the resulting parameter values. This interpretation step is complicated, however, by gauge freedoms— directions in parameter space along which model parameters can be changed without altering model predictions. When gauge freedoms are present in a model, the values of individual model parameters cannot be meaningfully interpreted without additional constraints. In standard Potts models of proteins, for example, the values of the parameters representing interactions between amino acids cannot be directly interpreted as quantifying interaction strength. This is because gauge freedoms make it possible to change any specific coupling parameter of interest without affecting model predictions by also making appropriate compensatory changes to other model parameters [2–6].

Researchers who study sequence-function relationships using quantitative models routinely encounter gauge freedoms. In practice, one of two methods is used to overcome the difficulties that gauge freedoms present. One method, called “gauge fixing”, removes gauge freedoms by introducing additional constraints on model parameters [2–18]. Another method limits the mathematical models that one uses to models that do not have any gauge freedoms in the first place [19–24]. Despite being frequently encountered in the course of research, the gauge freedoms present in models of sequence-function relationships have received only limited attention [e.g., 2, 4–6, 12, 25]. In particular, the mathematical properties of these gauge freedoms have yet to be systematically studied.

In physics, by contrast, gauge freedoms are well recognized as a topic of fundamental importance [26]. Gauge freedoms arise when a physical theory is expressed in a form that manifestly respects fundamental symmetries. For example, the classical theory of electricity and magnetism (E&M) is invariant to Lorentz transformations, i.e., relativistic changes in an observer’s velocity [27]. Lorentz invariance is obscured, however, when the equations of E&M are expressed directly in terms of electric and magnetic fields. To express these equations in a form that is manifestly Lorentz invariant, one must instead formulate them in terms of an electromagnetic four-potential. Doing this introduces gauge freedoms because the four-potential, unlike electric and magnetic fields, is neither directly measurable nor uniquely determined by the configuration of a physical system.[28] Nevertheless, working with the four-potential simplifies the equations of E&M and can aid in both their solution and their physical interpretation.

Motivated by the connection between gauge freedoms and symmetries in physics, we asked whether gauge freedoms in models of sequence-function relationships have a connection to the symmetries of sequence space, i.e., the possible ways of transforming the space of sequences without altering the Hamming distances between sequences. In this work we study the gauge freedoms of linear models that are equivariant under a specific symmetry group of sequence space—the group of position-specific character permutations (PSCP). Here, “linear models” are models that can be expressed as a sum of sequence features, each multiplied by a corresponding parameter; “PSCP” encompasses transformations that permute the identities of the individual characters (e.g., DNA bases or protein amino acids) at one sequence position, as well as transformations built from combinations of such permutations; and “equivariant” describes models for which linear transformations of the model parameters are able to compensate for the effects of PSCP transformations of sequences. Equivariant linear models include many of the most commonly used models in the literature, such as models with pairwise and/or higher-order interactions.

Using techniques from the theory of matrix representations of the symmetric group [29], we find that the gauge freedoms of these linear equivariant models arise when model parameters transform under redundant irreducible representations of PSCP. Based on this finding, we introduce an “embedding distillation” procedure that, for any linear equivariant model, facilitates both analytical and computational analyses of the vector space of gauge freedoms. We also study the connection between parameter interpretability and model transformation behavior. We find that in many (and possibly all) nontrivial models, the ability to interpret model parameters as quantifying the intrinsic effects of alleles requires that the model have gauge freedoms. This finding shows that models having gauge freedoms can have important advantages over models that have no gauge freedoms.

A companion paper [30] reports specific gauge-fixing strategies that can be applied to an important subset of the linear equivariant models, one that includes the most commonly used models of sequence-function relationships. It also describes specific ways of using these gauge-fixing strategies to assist in the development and biological interpretation of such models.

## BACKGROUND

We now establish definitions and notation used in Results. We also review basic results regarding gauge freedoms in mathematical models of sequence-function relationships. Our companion paper [30] provides an expanded discussion of these results together with corresponding proofs.

### Sequence-function relationships

Let *A* denote an alphabet comprising *α* distinct characters, let *S* denote the set of *α*^*L*^ sequences of length *L* built from these characters, and let *s*_*l*_ ∈ *A* denote the character at position *l* in any sequence *s* ∈ *S*. A real-valued model of a sequence-function relationship, 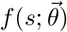,is defined to be a function that maps each sequence *s* to a real number. The vector 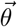 denotes the parameters of the model and is assumed to comprise *M* real numbers. For technical reasons it is sometimes useful to consider complex-valued models of sequence-function relationships, which are defined analogously.

### Linear models

Linear models of sequence-function relationships are linear in 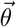 and thus have the form

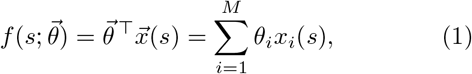

for all *s* ∈ *S*. Here, 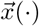 is an *M*-dimensional vector of sequence features, and each feature *x*_*i*_(·) is a function that maps *S* to ℝ. We refer to the space ℝ^*M*^ in which 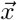 and 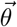 live as feature space.[31]

An example of a linear model is the pairwise one-hot model, which has the form

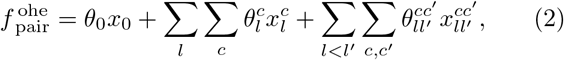

where the arguments of both the model and features have been kept implicit. In Eq. 2, *l, l*^′^ ∈ {1, …, *L*} index the positions within each sequence, and *c, c*^′^∈ *A* index the possible characters at these positions. We use the superscript “ohe” here and in what follows to indicate mathematical objects (such as embeddings, models, and representations) that are based on one-hot embeddings. Pairwise one-hot models, in particular, make use of the pairwise one-hot embedding 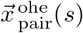,the elements of which represent three types of features: the constant feature, *x*_0_(*s*), which equals one for every sequence *s*; additive one-hot features, 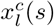,which equal one if *s*_*l*_ = *c* and equal zero otherwise; and pairwise one-hot features, 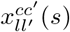,which equal one if both *s*_*l*_ = *c* and *s*_*l*_^*′*^ = *c*^′^, and equal zero otherwise.

### Gauge freedoms

Gauge freedoms are transformations of model parameters that do not affect model predictions. Formally, a gauge freedom is any vector 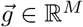 that satisfies

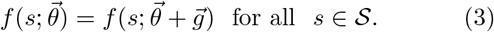

For linear sequence-function relationships the set of gauge freedoms, denoted by *G*, forms a vector space in ℝ^*M*^. It is readily shown that *G* is the orthogonal complement of the space spanned by sequence embeddings [30]. In what follows, we use *γ* to represent the dimension of *G*, i.e., the number of (independent) gauge freedoms.

Gauge freedoms arise from linear dependencies among sequence features. By inspection we see that 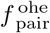 has

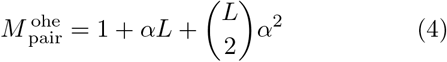

parameters. However, it turns out that the space spanned by the corresponding embedding 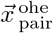 has only 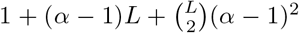 dimensions. This difference reflects the presence of 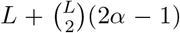 constraints on the features, namely: 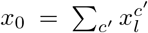 for all positions *l* (yielding 1 constraint per position); and 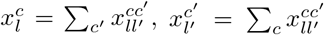 for all pairs of positions *l* < *l*^′^ and all choices of character *c* or *c*^′^ (yielding 2*α* − 1 independent constraints per pair of positions). The model 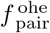 therefore has

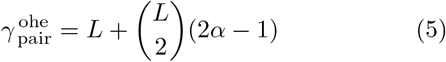

gauge freedoms. See our companion paper [30] for more details, as well as [2, 4, 6, 10] for earlier treatments of gauge freedoms in the pairwise one-hot model.

### Fixing the gauge

Fixing the gauge is the process of removing gauge freedoms by restricting 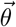 to a subset Θ of parameter space called “the gauge”. For example, the commonly used “zero-sum gauge” [4, 6] for the pairwise one-hot model is the subspace of parameter space in which the additive parameters at every position sum to zero when marginalized over characters (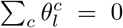 for every *l*) and the pairwise parameters at all pairs of positions sum to zero when marginalized over the characters at either position (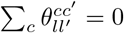 and 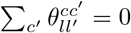 for every *l, l*^′^, *c, c*^′^).

Linear gauges are choices of Θ that are vector spaces.

The zero-sum gauge is one such linear gauge. A useful property of linear gauges is that gauge-fixing can be accomplished through linear projection. Specifically, for any linear gauge Θ, there exists a projection matrix *P* that projects each parameter vector 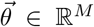 onto an equivalent parameter vector 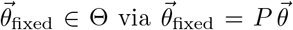.

Our companion paper describes a parametric family of linear gauges (including an explicit formula for the corresponding projection matrices) that includes as special cases many of the most commonly used gauges in the literature [30].

## RESULTS

We begin this section by formally defining the group of PSCP transformations, as well as the notion of model equivariance under this group. We then illustrate, for two example pairwise-interaction models, how transformation behavior under PSCP impacts both gauge freedoms and a specific type of parameter interpretability, namely the ability to assign intrinsic effects to individual alleles. Next we formally investigate this relationship more generally using methods from the theory of group representations [29]. In doing so, we establish an “embedding distillation” procedure that, for any equivariant model, enables analytic calculation of the number of gauge freedoms. We also establish an algorithm that enables the efficient computation of a sparse basis for the space of gauge freedoms. We conclude by revisiting the issue of parameter interpretability in light of these results.

### Position-specific character permutations (PSCP)

Different transformations of sequence space impact models of sequence-function relationships in different ways. Here we focus on PSCP transformations. These transformations of sequence space form a mathematical group, which we denote by *H*_PSCP_. The action of a transformation *H* ∈ *H*_PSCP_ on a sequence *s* ∈ *S* is written *Hs. H*_PSCP_ is a symmetry group of sequence space in that its transformations preserve the Hamming distances between sequences. There are other symmetry groups of sequence space as well, but we ultimately find that these symmetry groups do not have the same connections to gauge freedoms that *H*_PSCP_ does (discussed below and in SI Sec. 7).

### Equivariance

We also focus on equivariant linear models of sequence-function relationships. These are models for which both embeddings and parameters transform linearly under *H*_PSCP_. The specific sets of matrices that encode these linear transformations are called “representations” [29]. In general, a representation *R* of a group *H* is a function that maps each *H* ∈ *H* to a matrix *R*(*H*) in a way that preserves the multiplicative structure of *H*, i.e., *R*(*H*_1_*H*_2_) = *R*(*H*_1_)*R*(*H*_2_) for any two group elements *H*_1_, *H*_2_ ∈ *H*. The degree of the representation *R* (denoted deg *R*) is the dimension of the vector space on which *R* acts. Two different examples of representations for the same group are described and illustrated below (see Eqs. 8 and 11, as well as Fig. 1)

**Figure 1.**
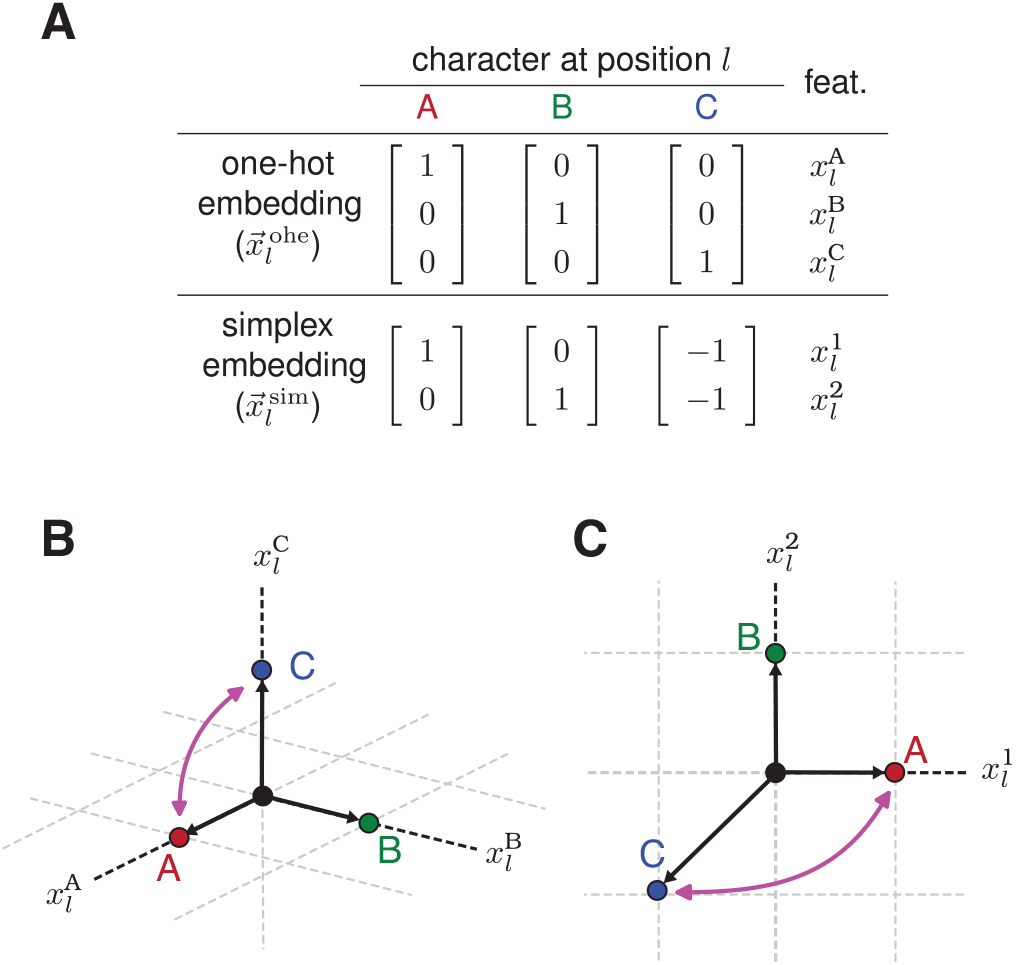
Transformation behavior of two single-position embeddings. (A) Two possible embeddings of characters at position *l* of a sequence built from the three-character alphabet *A*= {A, B, C}: the three-dimensional one-hot embedding 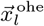 and the two-dimensional simplex embedding 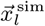 The elements of 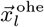 are the three one-hot sequence features 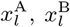,and 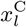.The two elements of 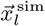 are denoted 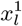 and 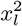. (B) The three-dimensional embedding 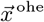 for each possible character at position *l*. (C) The two-dimensional embedding 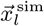 for each possible character at position *l*. Pink arrows in panels B and C indicate the transformation of each embedding vector induced by permuting the characters A and C.

Formally, we say that an embedding 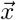 is equivariant in *H* if and only if there is a representation *R* of *H* such that

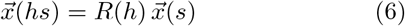

for all *H* ∈ *H* and all ∈ *s S*. We also say that a model is equivariant if and only if it has an equivariant embedding. For an equivariant model whose embedding transforms as in Eq. 6, the transformation of *S* by any *H* ∈ *H* can be compensated for by the transformation of 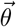 by *R*(*H*)^−1⊤^, in the sense that

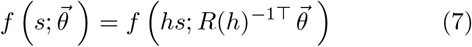

for every *s* ∈ *S* and every 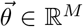 (see SI Sec. 3.2). Although linear models of sequence-function relationships can be equivariant in a variety of symmetry groups *H*, we use the term “equivariant” to specifically refer to equivariance under *H*_PSCP_ unless otherwise noted.

### One-hot models

The most commonly used equivariant models are based on single-position one-hot embeddings. Such models are arguably the most intuitive, as their features are built from the indicator functions for single-position alleles (e.g., the nucleotides in a DNA sequence or the amino acids in a protein sequence). We denote the single-position one-hot embedding for position *l* as 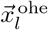 And define it to be a binary vector of dimension *α* with features 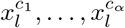,where *c*_1_, …, *c*_*α*_ is an ordering of the characters in *A*. For example, Fig. 1A shows 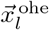 for the three-character alphabet *A* = {A, B, C}.

The embedding 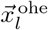 transforms under what is known as a “permutation representation” [29]. We denote this representation as 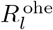.For example, consider the transformation *H*_A↔C_ that exchanges characters *A* and *C* at every position in a sequence. The effect of this transformation on 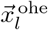 (Fig. 1B) is equivalent to multiplying 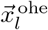 by the matrix

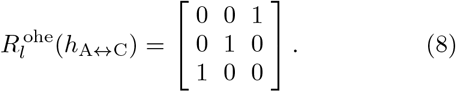

This and all other matrices in the representation 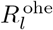 are permutation matrices, in that all matrix elements are 0 or 1, and each row and column contains a single 1. Consequently, multiplying a vector by one of these matrices changes the order of the elements in that vector, but does not change the overall set values that those elements take. We refer to 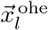 and other embeddings that transform under permutation representations as permutation embeddings; their corresponding models are called permutation models.

The embeddings of many different models can be built by taking direct sums of Kronecker products 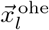. For example, the pairwise one-hot model of Eq. 2 is based on the embedding

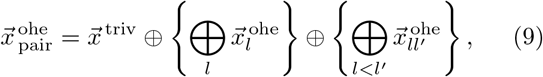

Where 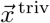 denotes the trivial embedding (defined to be the 1-dimensional vector [1] for all sequences) and the Kronecker product

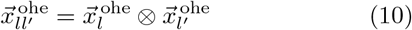

yields an *α*^2^-dimensional embedding having elements 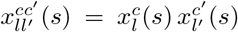 for all *s* ∈ *S* and all characters *c, c*^′^ ∈ *A*. The direct sums in Eq. 9 yield 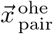 by stacking the component embeddings on top of one another in the resulting column vector. Note that, because 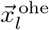 is a permutation embedding, so is 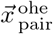. In fact, any embedding constructed from a direct sum of Kronecker products of 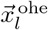 is a permutation embedding. We call this class of models the “generalized one-hot models”.

How a single-position embedding transforms has important consequences for how the parameters of models constructed from that embedding are interpreted. For the pairwise one-hot model, the fact that 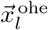 transforms under a permutation representation implies that both 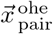 and 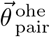 transform under permutation representations as well. A consequence of this is that the individual parameters in 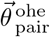 can be interpreted as quantifying the intrinsic effects of individual alleles. For example, the transformation *H*_A↔C_ induces a permutation of parameters that exchanges 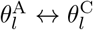 at all positions *l*, exchanges 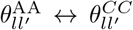 at all pairs of positions *l* < *l*^′^, and so on.

Model parameters therefore track their corresponding alleles: 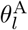 tracks sequences that have A at position 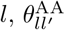 tracks sequences that have AA at positions *l* and *l*^′^, etc.. The fact that 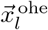 transforms under a permutation representation also means that the features therein are not linearly independent. For example, the three embedding vectors in Fig. 1B lie within a two-dimensional affine subspace defined by the constraint 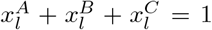 As we will see, a consequence of such constraints is that embeddings (like 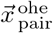) that are built from direct sums of Kronecker products of single-position one-hot embeddings will yield models that have gauge freedoms. So although the parameters of generalized one-hot models can be interpreted as quantifying intrinsic allelic effects, the numerical values of individual parameters cannot (at least in some cases) be interpreted in the absence of gauge-fixing constraints.

### Simplex models

Simplex embeddings mathematically represent alleles in a more compact but less intuitive way than the one-hot embeddings discussed above do. Single-position simplex embeddings encode the *α* characters of *A* using zero-centered vectors of dimension *α* − 1, and thus have fewer dimensions than corresponding alleles. Simpilex embeddings can be defined in multiple ways that differ from one another by similarity transformations, i.e., change-of-basis transformations. Here we adopt a particularly convenient definition: 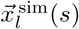 is defined to be an *α* − 1 dimensional vector, the *i*’th element of which is 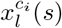 if *s*_*l*_ ≠ *c*_*α*_ and −1 if *s*_*l*_ = *c*_*α*_. We use the superscript “sim” here and in what follows to indicate mathematical objects that are based on simplex embeddings. Figs. 1A,C illustrate 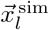 for the three-character alphabet. Unlike 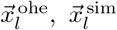 transforms under a non-permutation representation, which we denote as 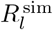 For example, the effect of *H*_A↔C_ on 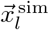 is equivalent to multiplication by the matrix

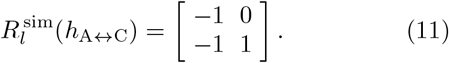

As with one-hot embeddings, the embeddings of many different models can be built from direct sums of direct products of 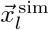. For example, a simplex embedding analogous to 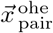 can be constructed as

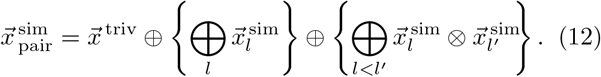

The corresponding pairwise simplex model has the form

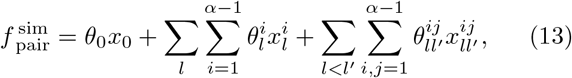

where 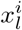 denotes the *i*’th element of 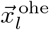, and where 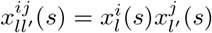 for all *s* ∈ *S*. We use 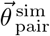 denote the parameters of this model. Note that these parameters are indexed using numerical superscripts ranging from 1 to *α*− 1, rather than by characters in *A*.

Pairwise simplex models describe the same sequence-function relationships that pairwise one-hot models do, i.e., given a set of parameters for one of these models, there exists a corresponding set of parameters for the other model that yields the same predictions over all sequences. However, because 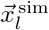 has lower dimension than 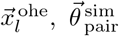 contains fewer parameters than 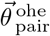. Inspection of Eq. 12 shows that the number of parameters in 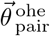 is, in fact,

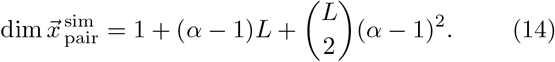

This reduction in the number of parameters entirely eliminates gauge freedoms, as can be seen from the fact that

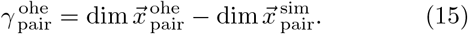

The lack of gauge freedoms in 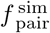 is one example of the fact that, as we will see, models defined using (non-redundant) simplex embeddings do not have gauge freedoms. In fact, multiple groups [20, 22, 23] have argued for the use of simplex models, rather than one-hot models, based on the simplex models not having gauge freedoms.

We argue, however, that the parameters of simplex models are fundamentally more difficult to interpret as allelic effects than are the parameters of one-hot models. Because 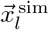 does not transform under a permutation representation, neither does 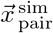 and neither does 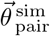. In the case of the three-character alphabet, one sees from Eq. 12 that *H*_A↔C_ induces a transformation of model parameters that maps 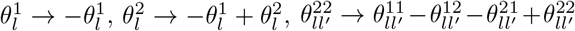, and so on. The fact that these parameters change in ways described by nontrivial linear combinations means that individual parameters cannot be interpreted as quantifying individual allelic effects.

### Maschke decomposition

We now use methods from the theory of group representations to formally investigate the general connection between model transformation behavior and gauge freedoms. Maschke’s theorem, a foundational result in representation theory, states that every matrix representation of a finite group is equivalent to a direct sum of irreducible matrix representations. Here the term “equivalent” means that there is a similarity transformation (i.e., a change of basis) that maps one representation to another; we use the symbol ≃ to denote equivalence in what follows. The term “irreducible” means that the representation has no proper invariant subspace.

Consider for example 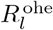,the representation that describes how 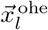 transforms under *H*_PSCP_. The group *H*_PSCP_ is isomorphic to the symmetric group (i.e., the group of permutations), the representations of which are well understood [29]. In this context, 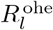 is called the “defining representation” and is well known to be reducicble. Specifically, 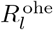 has two proper invariant subspaces. One subspace has dimension 1 and is spanned by the vector [1 1 · · · 1]^⊤^. The other subspace has dimension *α* − 1 and consists of the set of *α*-dimensional vectors whose elements sum to zero. The first of these subspaces transforms under the “trivial representation”, which is simply the 1×1 matrix [1] and which we denote by 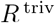.The other subspace transforms (after an appropriate change of coordinates) under the representation 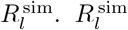 is called the “standard representation” and is well known to be irreducible. The Maschke decomposition of 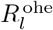 is therefore given by

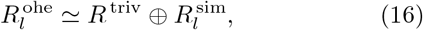

where the direct sum on the right-hand side yields a block diagonal matrix created from *R*^triv^and 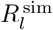.

Equivalently, we can think of Maschke decomposition in terms of embeddings. Thinking in terms of embeddings can be helpful for deriving the specific invertible matrix that performs the similarity transformation needed to express a Maschke decomposition as an equality instead of an equivalence. When multiplied by an appropriate similarity transformation matrix 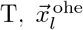 can be expressed as a direct sum of the trivial embedding 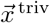 (which is simply the 1-dimensional vector [1]) and the simplex embedding 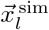, i.e.,

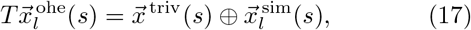

for all sequences *s*. This allows us to express the equivalence relation in Eq. 16 as an equality, as it implies that

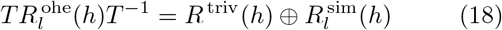

for all group elements *H*. Based on the definition of the embeddings 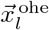 and 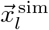 above, one can readily show that the similarity transformation matrix *T* is given by

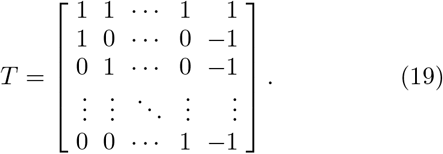

This matrix *T* will be used later when defining an algorithm for distilling general equivariant embeddings.

### Decomposition of equivariant embeddings

Maschke’s Theorem implies that any representation *R* of *H*_PSCP_ can be expressed as

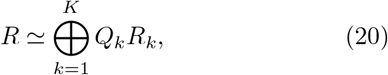

where the *R*_*k*_ are distinct irreducible representations of *H*_PSCP_ and each *Q*_*k*_ is a natural number that denotes the multiplicity of *R*_*k*_ in the direct sum. *R* is thus equivalent to a block-diagonal representation formed by placing *Q*_*k*_ copies of each *R*_*k*_ along the diagonal and setting all other matrix elements to zero (see Fig. 2B). One consequence of Eq. 20 is that any embedding 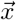 that transforms under *R* can be decomposed as

**Figure 2.**
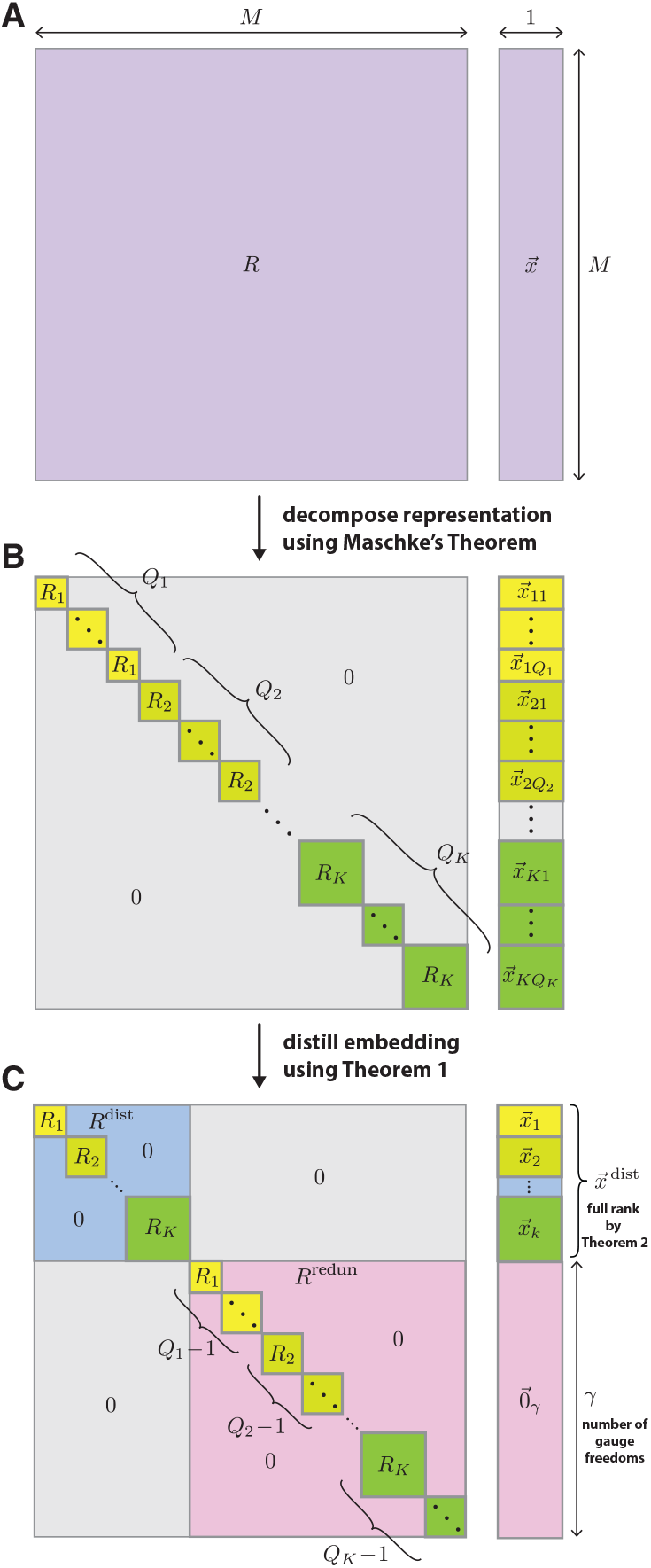
Embedding distillation. (A) Given an *M*-dimensional embedding 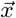 that is equivariant under *H*_PSCP_, let *R* be the representation that describes how 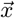 transforms. (B) By Maschke’s theorem, *R* can be decomposed into a direct sum of irreducible representations, *R*_*k*_ (*k* ∈{1, …, *K*}), each of which occurs with multiplicity *Q*_*k*_ (Eq. 20). Similarly, 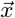 can be decomposed into a direct sum of irreducible embeddings 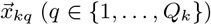, where each 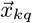 transforms under *R*_*k*_ (Eq. 21). (C) By Theorem 1, an additional similarity transformation can be performed that, for each value of *k*, zeroes out all but one 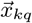 and sorts the remaining embeddings; each remaining 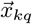 is denoted by 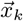. Consequently,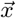 decomposes into a direct sum of a distilled embedding 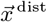 and a zero vector 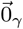 having some dimension *γ* (Eq. 24). 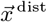 is given by the direct sum of all 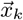 (Eq. 25) and is full rank by Theorem 2. The distilled representation *R* ^dist^ describes how 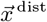 transforms and is given by a direct sum of one copy of each *R*_*k*_. The redundant representation *R* ^redun^ operates On 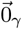 and comprises the *Q*_*k*_− 1 remaining copies of each *R*_*k*_. The resulting number of gauge freedoms is *γ* (see Eq. 28).

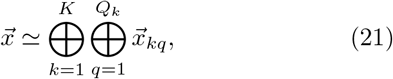

where each 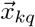 is an embedding that transforms under *R*_*k*_. In what follows, we say that embeddings like 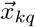 are irreducible because they transform under irreducible representations. We also assume that all 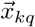 are nonzero, but this assumption can be removed without fundamentally changing our results; see SI Sec. 5.2 for details. The Maschke decompositions of *R* and 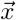 are illustrated in Fig. 2A,B.

### Distillation of equivariant embeddings

We now describe an “embedding distillation” procedure that connects the Maschke decomposition of 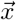 to the gauge freedoms of the corresponding model. In SI Sec. 5.1 we prove the following:

#### Theorem 1

*Any two nonzero sequence embeddings that transform under the same irreducible representation of H*_PSCP_ *are equal up to a constant of proportionality*.

Using Theorem 1 we find that there is a similarity transformation matrix *T*_decom_ such that

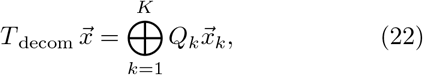

where, for each *k*, 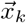 denotes any one of the irreducible embeddings 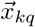 in Eq. 21 and *Q*_*k*_ denotes the multiplicity of each term in the direct sum. Next we perform a similarity transformation (described by a matrix *T*_thin_) that “thins out” the embedding by setting all except one copy of each 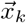 to zero. Finally, we perform a similarity transformation (described a matrix *T*_sort_) that “sorts” the remaining nonzero embeddings, arranging them in series at the top of the resulting embedding vector. We thus find that applying the cumulative similarity transformation given by

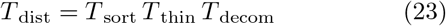

to the embedding 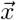 yields

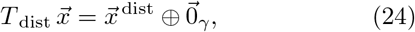

Where 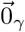 is a *γ*-dimensional vector of zeros and

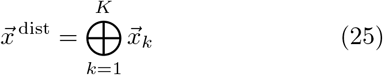

is a “distilled embedding”. When applied to the representation *R*, this distillation procedure yields

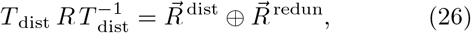

where the “distilled representation”, 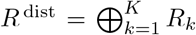, comprises one copy of each *R*_*k*_ present in Eq. 20, and where the redundant representation, 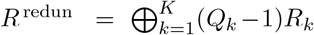 sweeps up the remaining copies of each *R*_*k*_. The final distilled versions of *R* and 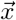 are illustrated in Fig. 2C. Explicit formulae for constructing *T*_decom_, *T*_thin_, and *T*_dist_ are given in SI Sec. 8.

### Identification of gauge freedoms in equivariant models

To identify the gauge freedoms of an equivariant model, we use the fact that 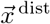 (defined in Eq. 25) is full rank. This is a consequence of the following Theorem, which is proven in SI Sec. 3.4:

#### Theorem 2

*For each k* ∈{1, …, *K*}, *let* 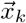 *be a nonzero embedding that transforms under an irreducible representation R*_*k*_ *of the group H*_PSCP_. *Then the direct sum of all* 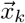 *is full rank if and only if all R*_*k*_ *are pairwise inequivalent*.

Because 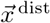 is full rank, 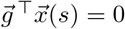 for all *s* ∈*S* if and only if

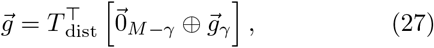

where *T*_dist_ is the distillation matrix in Eq. 23 and 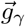 is any vector in ℝ^*γ*^. The space of gauge transformations *G* is therefore given by the set of vectors 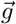 that have the form in Eq. 27. In particular, the number of gauge freedoms is seen to be

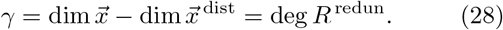

We thus see that the number of gauge freedoms of an equivariant linear model is equal to the sum of the degrees of all the redundant irreducible representations under which that model’s embedding (or equivalently, that model’s parameter vector) transforms.

### Identification of all equivariant models

The mathematical structure of a group defines the models that transform equivariantly under that group. In the case of *H*_PSCP_, the relatively simple group structure allows the straight-forward identification of all inequivalent distilled embeddings and thus the identification of all equivariant linear models.

*H*_PSCP_ can be written as a direct product of simpler groups:

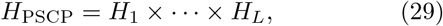

where each *H*_*l*_ denotes the group of character permutations at sequence position *l*. Each irreducible representation *R*_*k*_ of *H*_PSCP_ can therefore be expressed as the

Kronecker product

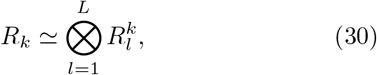

where each 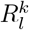 is an irreducible representation of *H*_*l*_ (see Theorem 1.11.3 of [29]). An embedding 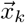 that trans-forms under *R*_*k*_ will therefore have the form

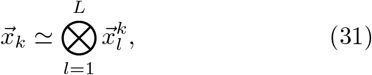

where 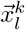 is an irreducible embedding that transforms under *H*_*l*_. In SI Sec. 4.3 we show that *H*_*l*_ supports only two inequivalent irreducible embeddings (regardless of alphabet size): 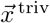 and 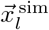, Each 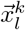 must therefore be equivalent to one of these two embeddings. Ignoring the factors of 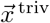 because they do not impact Kronecker products, Eq. 31 becomes

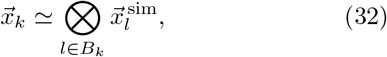

where *B*_*k*_ is a subset of the positions {1, …, *L*}. There are 2^*L*^ possible choices for each subset *B*_*k*_, and thus 2^*L*^ inequivalent irreducible embeddings 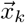. Since each 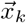 can appear at most once on the left-hand side of Eq. 25, there are 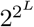 inequivalent distilled embeddings 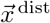.

For each choice of 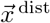 there are an infinite number of possible choices for *T*_dist_ and *γ* that can be used to define 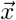 (via Eq. 24). The number of possible equivariant embeddings 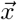, and thus the number of equivariant models *f*, is therefore infinite. However, all models corresponding to a specific 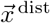 have the same expressivity, i.e., the set of sequence-function relationships that each model describes (considered over all possible values of model parameters) is the same. We therefore consider these models to be equivalent, and conclude that there are a total of 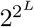 inequivalent equivariant linear models on sequences of length *L*.

### Analytical analysis of generalized one-hot models

We now use the embedding distillation procedure to compute the number of gauge freedoms of all generalized one-hot models. This derivation is based on the Maschke decomposition 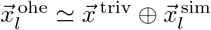 from Eq. 17.

We first demonstrate this calculation on the pairwise one-hot model. Plugging the decomposition of 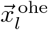 into the definition for 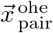 in Eq. 9, then expanding the Kronecker products and grouping like terms, we find that

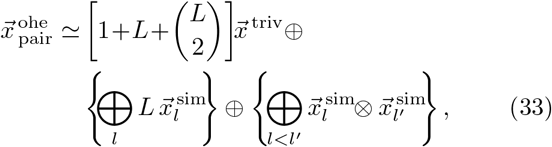

where the scalar coefficients correspond to the *Q*_*k*_ in Eq. 22. We derive the corresponding distilled embedding by replacing each of these coefficients with 1. Doing so reveals the distillation of 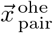 to be 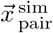. The result for 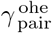 in Eq. 5 is therefore just a manifestation of Eq. 28. We now extend this approach to all generalized one-hot models. The embedding 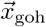 of any generalized one-hot model can be written as

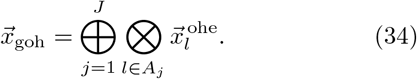

where *A*_1_, …, *A*_*J*_ denote *J* (not necessarily distinct) sets of positions. Because the dimension of 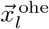 is *α*, the number of corresponding model parameters is

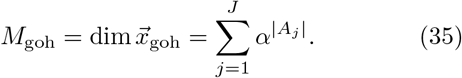

Decomposing 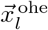 in terms of 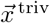 and 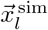, expanding each Kronecker product, then grouping the resulting terms, we find that

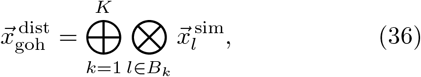

where *B*_1_, …, *B*_*K*_ denote the distinct subsets of positions that occur among all the *A*_*j*_. Because the dimension of 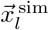 is *α* − 1,

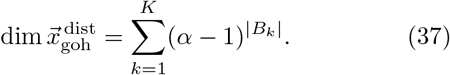

The number of gauge freedoms of the generalized one-hot model having embedding 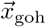 is therefore given by

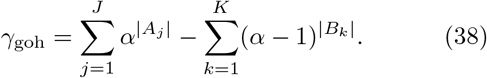

Table I reports the number of gauge freedoms calculated in this manner for a variety of generalized one-hot models (illustrated in Fig. 3). SI Sec. 6 provides expanded descriptions for each generalized one-hot model, as well as detailed derivations of the results in Table I.

**Table I.**
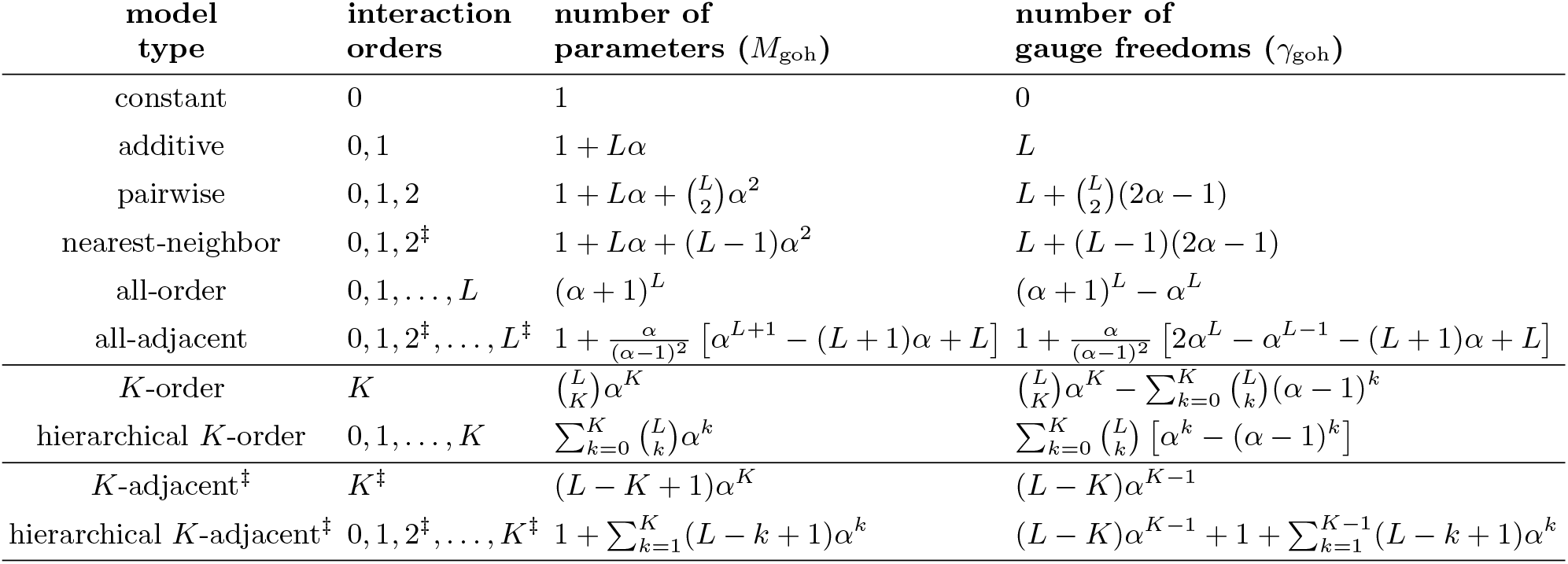
Analytical results for various generalized one-hot models, computed using Eqs. 35 and 38. See SI Sec. 6 for derivations of these results. *K*-adjacent models assume *K* ≥ 1. ^‡^Only includes interactions among adjacent positions.

**Figure 3.**
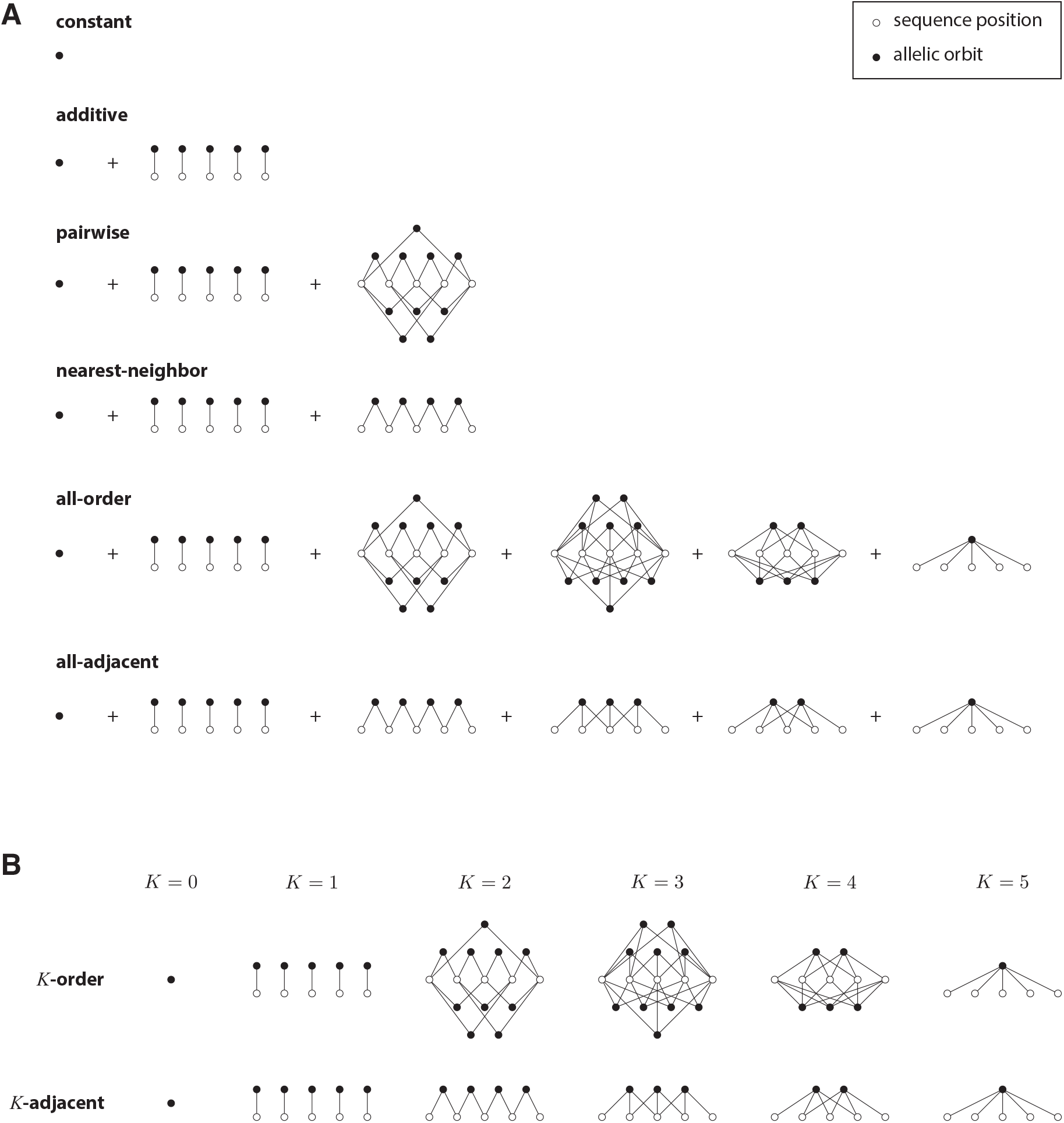
Structure of generalized one-hot models analyzed in Table I for sequences of length *L* = 5. Open circles represent sequence positions. Closed circles represent allelic orbits, i.e., sets of sequence features that are closed under the action of *H*_PSCP_. Edges indicate position indices shared by the features in each allelic orbit. (A) Structure of constant, additive, pairwise, nearest-neighbor, all-order, and all-adjacent models. (B) Structure of *K*-order models and *K*-adjacent models for various interaction orders *K*.

A result of this analysis is that all generalized one-hot models have gauge freedoms, save models for which the direct sum in Eq. 34 includes only one term. To see this, observe that Eq. 22 gives

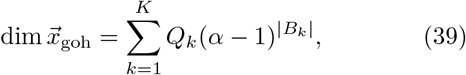

where each multiplicity value *Q*_*k*_ is equal to the number of sets *A*_*j*_ that contain *B*_*k*_. Using this together with Eq. 36 and Eq. 28 gives

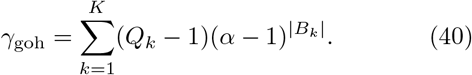

We thus find that *γ*_goh_ = 0 if and only if none of the *Q*_*k*_ are greater than 1. But since the empty set is a subset of every *A*_*j*_, it will always be among the *B*_*k*_, and the corresponding multiplicity value will be *Q*_*k*_ = *J*. Gauge freedoms are therefore present in all generalized one-hot models for which *J* ≥ 2. Conversely, *γ*_goh_ = 0 when *J* = 1 because all *B*_*k*_ occur with multiplicity *Q*_*k*_ = 1. Gauge freedoms are therefore absent in all generalized one-hot models for which *J* = 1.

### Computational analysis of generalized one-hot models

Embedding distillation also allows one to efficiently compute a sparse basis for the space of gauge freedoms *G*_goh_ of any generalized one-hot model. Eq. 27 reveals that *G*_goh_ is spanned by the last *γ*_goh_ row vectors of *T*_dist_. One can therefore compute a basis for *G*_goh_ by computing *T*_dist_. This is notable because computing *T*_dist_ only requires keeping track of the similarity transformations needed to express 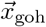 in the distilled form shown in Eq. This computation is far less demanding than computing a basis for *G*_goh_ using Gaussian elimination or singular value decomposition when (as is often the case) the number of possible sequences is far greater than the number of model parameters.

In Eq. 23 we described how to construct *T*_dist_ from a product of three matrices: *T*_decom_, *T*_thin_, and *T*_sort_. Explicit formulae for computing these matrices, as well as their inverses, are provided in SI Sec. 8. For these formulae we observe that each matrix, as well as its inverse, is sparse in the large *L* limit when the maximal order of interaction described by the model is fixed. The resulting distillation matrix *T*_dist_ is therefore also sparse, as is its inverse. It also turns out that every nonzero element of *T*_dist_ is +1 or −1. Taking the last *γ*_goh_ rows of *T*_dist_ thus provides a basis for *G*_goh_ consisting of sparse vectors whose only nonzero elements are +1 and −1. Having sparse matrices for *T*_dist_ and 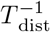 also allows us to compute a sparse gauge-fixing projection matrix *P*; see SI Sec. 8 for details. Fig. 4 illustrates the actions of *T*_decom_, *T*_thin_, and *T*_sort_ on an example embedding vector for the all-order interaction model corresponding to *L* = 3 and *α* = 3. Fig. 4 also illustrates the corresponding distillation matrix *T*_dist_.

**Figure 4.**
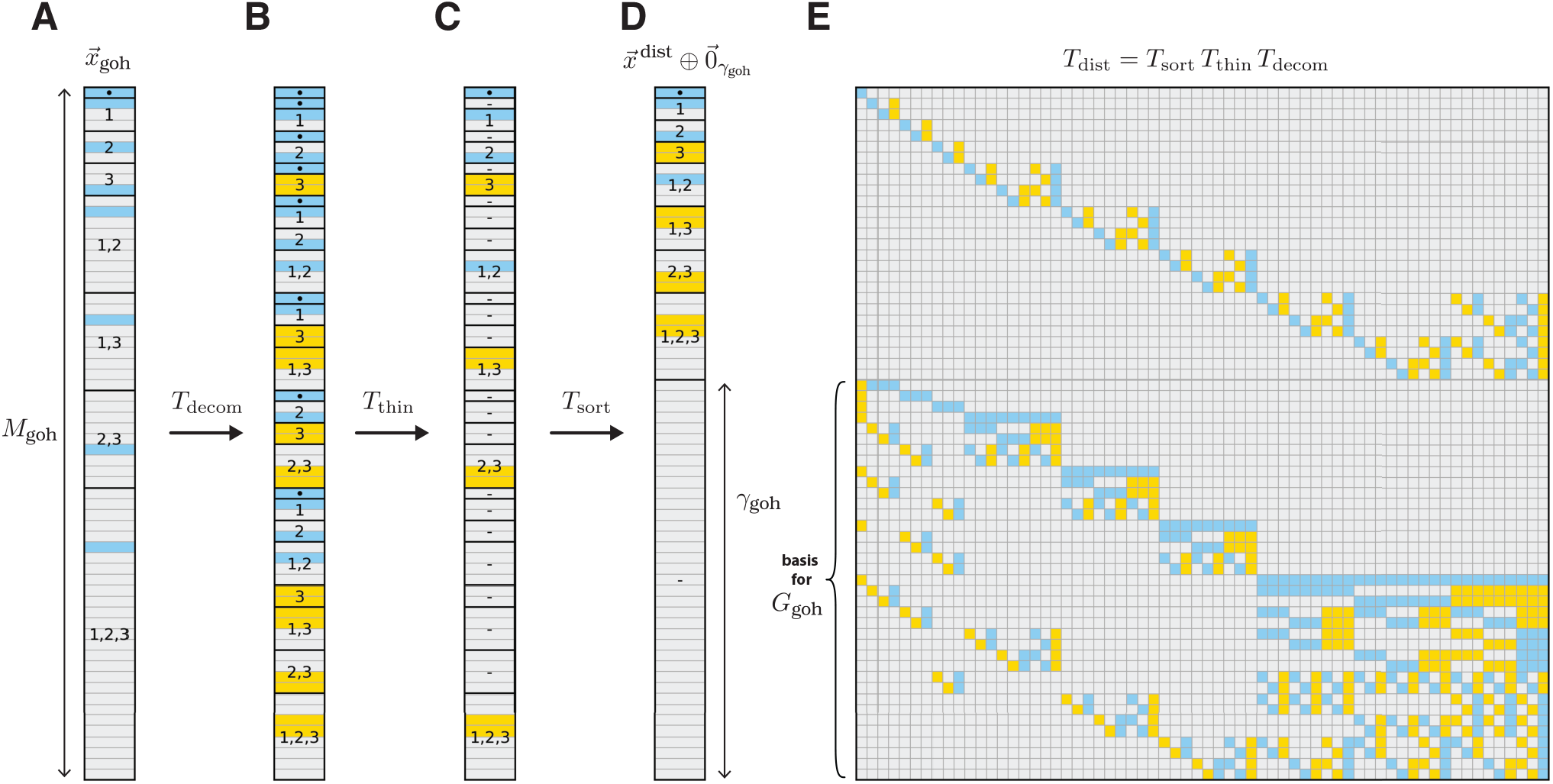
Embedding distillation for an example generalized one-hot model. (A) Embedding 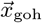 of the *L* = 3 sequence *s* = ABC for an all-order one-hot model based on the alphabet *A*= {A, B, C}. This embedding has degree *M*_goh_ = 64. (B) Result of multiplication by the decomposition matrix *T*_decom_. (C) Result of subsequent multiplication by the thinning matrix *T*_thin_. (D) Result of subsequent multiplication by the sorting matrix *T*_sort_, which yields 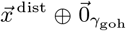 with *γ*_goh_ = 37 being the number of gauge freedoms. In B-D, dots indicate 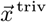, dashes indicate zero vectors, and numbers indicate the positions *l* contributing to each component. (E) Distillation matrix *T*_dist_ that implements the full distillation procedure in A-D. The last *γ*_goh_ rows of *T*_dist_ provide a sparse basis for the gauge space, *G*_goh_. In A-E, vector and matrix elements are colored according to their numerical values: blue represents +1, yellow represents −1, and gray represents 0.

### Other symmetry groups

The proof of Theorem 1 in SI Sec. 5.1, and thus our embedding distillation procedure, applies only to the symmetry group *H*_PSCP_. There are other symmetry groups of sequence space besides *H*_PSCP_, however, and it is worth asking whether Theorem 1, and thus Eqs. 24-28, hold for those groups as well.

One symmetry group is the group of global character permutations, *H*_GCP_. This group comprises transformations that apply the same permutation to characters at every position in a sequence. Another is the group of position permutations, *H*_PP_. This group comprises transformations that permute the positions in a sequence without otherwise changing the characters therein. SI Sec. 7.1 shows that Theorem 1 does not hold for either *H*_GCP_ or *H*_PP_. Consequently, one cannot compute distilled embeddings using the irreducible representations of either group.

A third symmetry group is *H*_Ham_, which describes combinations of position permutations and position-specific character permutations. *H*_Ham_ is the largest symmetry group that preserves Hamming distances [32], and includes *H*_PSCP_, *H*_PP_, and *H*_GCP_ as subgroups. Theorem 1 does hold for *H*_Ham_, due the fact that *H*_PSCP_ is a subgroup (see SI Sec. 7.2). However, the set of models that are equivariant under *H*_Ham_ is a subset of the models that are equivariant under *H*_PSCP_, and the irreducible representations of *H*_Ham_ are more complex than those of *H*_PSCP_. *H*_PSCP_ is therefore more useful than *H*_Ham_ for analyzing gauge freedoms.

### Transformation behavior and parameter interpretability

We now return to the connection between parameter transformation behavior and parameter interpretability. Our above discussion of pairwise models suggested that the ability to interpret individual parameters as quantifying intrinsic allelic effects required the presence of gauge freedoms. We now formalize this observation and conjecture an extension to all linear equivariant models.

Mathematically, we define a generalized allele *A* to be any subset of *S*, and say that any sequence *s* ∈ *S* has allele *A* if *s* ∈*A*. The corresponding “allelic feature” *x*_*a*_ is defined to be the indicator function on *S* for whether a sequence has allele *A*. An “allelic model” is defined to be a linear sequence-function relationship in which every feature is an allelic feature. In the context of an allelic model, the parameter θ_*a*_ that multiplies *x*_*a*_ is said to be an “allelic effect.” The parameters of a linear model can therefore be interpreted as allelic effects if and only if every one of the corresponding features is an indicator function for membership in some subset of *S*.

For an allelic model to have parameters that describe intrinsic allelic effects, the model must be a “permutation model”, i.e., the features and parameters of the model must transform under a permutation representation of *H*_PSCP_. Requiring an allelic model to be a permutation model puts strong constraints on which sets of alleles it can describe. Because *H*_PSCP_ permutes sequences, it also permutes alleles. Given a specific allele *A*, we call the set of alleles created by the action of *H*_PSCP_ on *A* an “allelic orbit”. It is readily seen that, for an allelic model to be a permutation model, the set of alleles it describes must consist of some number *J* of complete allelic orbits.

All allelic models that comprise *J* ≥ 2 allelic orbits have gauge freedoms. To see this, observe that the features in each orbit transform among themselves according to a permutation representation. The features of the full model will therefore transform under a direct sum of *J* permutation representations. Because every permutation representation contains the trivial representation in its Maschke decomposition, the decomposition of the full model’s representation will contain at least *J* copies of the trivial representation. The model will therefore have at least *J* − 1 gauge freedoms, though additional gauge freedoms can be present as well.

This result is reflected in our above analytic analysis of generalized one-hot models. All generalized one-hot models are allelic permutation models (though the converse is not true; see SI Sec. 9.1), and each allelic orbit of a generalized one-hot model corresponds to a position set *A*_*j*_ in Eq. 34. The lower-bound on the number of gauge freedoms identified here recapitulates the finding above that generalized one-hot models have no gauge freedoms if and only if *J* = 1.

An allelic permutation model that does not have gauge freedoms must therefore comprise only one allelic orbit. An example of a model with only one allelic orbit is a one-hot model of length *L* = 1, e.g., a model describing the effects of only one nucleotide position in a DNA sequence or one amino acid position in a protein sequence. Are single-orbit allelic models useful in practice? We argue that the answer is essentially no. In SI Sec. 9.1 we show that single-orbit generalized one-hot models cannot describe co-occurring alleles. We regard such models as trivial because the entire reason for quantitatively modeling sequence-function relationships is to deconvolve the influence of co-occurring alleles. There are single-orbit allelic permutation modelsthat describe co-occurring alleles, but all the examples of these we have analyzed either have gauge freedoms or are mathematically equivalent to generalized one-hot models (see SI Sec. 9.1). Moreover, among models whose embeddings are built from direct sums of Kronecker products of single-position embeddings, the generalized one-hot models have the fewest gauge freedoms (see SI Sec. 9.2). Based on these findings, we conjecture that all nontrivial allelic permutation models (i.e., all models whose parameters describe intrinsic allelic effects) have gauge freedoms.

## DISCUSSION

Motivated by the connection between gauge freedoms and symmetries in physics, we investigated the relationship between gauge freedoms and symmetries in quantitative models of sequence-function relationships. We found that, for linear models that are equivariant under *H*_PSCP_ (i.e., the group of PSCP transformations), gauge freedoms arise due to model parameters transforming under redundant irreducible matrix representations. From a conceptual standpoint, this result links the gauge freedoms of models of sequence-function relationships to the transformation behavior of these models under a specific symmetry group of sequence space. From a practical standpoint, this result facilitates the analytic calculation of the number of independent gauge freedoms in a large class of commonly used models. It also enables an embedding distillation algorithm that can efficiently compute a sparse basis for the space of gauge freedoms. This latter capability may prove to be useful particularly when studying models with very large numbers of parameters. Such models are increasingly common, as massively parallel reporter assays, deep mutational scanning experiments, and other multiplex assays of variant effect can now readily measure the activities of hundreds of thousands of sequences in a single experiment [e.g. 33, 34].

We also investigated the link between parameter transformation behavior and parameter interpretability. In doing so, we identified an incompatibility between two different notions of parameter interpretability: in linear models that are equivariant under *H*_PSCP_, the ability to interpret individual parameters as quantifying intrinsic allelic effects requires that these parameters transform under a permutation representation of *H*_PSCP_. But in many (and possibly in all) nontrivial models, this requirement is incompatible with the ability to interpret the values of individual parameters in the absence of gauge-fixing constraints. Consequently, models that have gauge freedoms can have advantages over equally expressive models that do not have gauge freedoms.

It should be noted that there are indeed useful models that do not have gauge freedoms. One such class of models are the “wild-type” one-hot models, the features of which are limited to those describing mutations away from a specific sequence of interest [e.g., 34, 35].

Note that wild-type models differ in an important way from one-hot models expressed in the wild-type gauge (described in [30]): the latter models have specific parameters set to zero, whereas the former models lack these parameters entirely.

The parameters of wild-type models have a close connection to the quantities that one can actually experimentally measure, i.e., activity differences between alleles. However, these parameters do not transform under a permutation representation of *H*_PSCP_ and so do not quantify intrinsic allelic effects. Indeed, wild-type models are quite close in spirit to the representation of E&M explicitly in terms of electric and magnetic fields: while these fields are the directly measurable manifestation of E&M, they transform in complicated ways under changes in velocity and so do not provide the theoretical clarity– the *intrinsic* description of E&M–that the electromagnetic four-potential does.

Another class of useful models that do not have gauge freedoms are models whose features represent sequence-dependent physical properties, such as the chemical properties of amino acids [36, 37] or the physical shape of the DNA double helix [38, 39]. These models are not equivariant, however, and their parameters describe the effects of physical properties of alleles, not the effects of alleles themselves. Notably, both classes of model reflect inductive biases that break *H*_PSCP_ symmetry.

In classical field theories like E&M, there are specific symmetries that are well-established by experiment and that any mathematical formulation of the theory must be consistent with. This does not, however, mean that the equations of the theory must transform in a simple way under those symmetries. Mathematically formulating physical theories so that the equations themselves manifestly respect the symmetries of the theory generally requires over-parameterizing the equations, thereby introducing gauge freedoms. Physicists often find it worthwhile to do this, as having fundamental symmetries be manifestly reflected in one’s equations can greatly facilitate the interpretation and application of those equations. Solving such equations, however, requires fixing the gauge—introducing additional constraints that make the solution of the equations unique.

Unlike in physics, there is no experimentally established requirement that models of sequence-function relationships be equivariant under any symmetries of sequence space. The specific mathematical form one uses for such models is subjective, and different models are commonly used in different contexts. Citing the ambiguities caused by gauge freedoms, some have argued for restricting one’s choice of model to those that have no gauge freedoms. Nevertheless, models that have gauge freedoms are still common in the literature. We suggest that a major reason for this may be that researchers often prefer to use models that manifestly reflect symmetries of sequence space, and therefore have parameters that are interpretable as intrinsic allelic effects. As we showed, these criteria often (and possibly in all nontrivial cases) require the use of over-parameterized models. In this way, the origin of gauge freedoms in models of sequence-function relationships does mirror the origin of gauge freedoms in physical theories.

There is still much to understand about the relationship between models of sequence-function relationships, the symmetries of these models, and how these models can be biologically interpreted. This paper and its companion [30] have only addressed gauge freedoms and symmetries in linear models. Some work has explored the gauge freedoms and symmetries of nonlinear models of sequence-function relationships [40, 41], but important questions remain. The sloppy modes [42, 43] present in these models are also important to understand, and to our knowledge these have yet to be systematically investigated. Addressing these problems is becoming increasingly urgent due to the expanding interest in interpretable quantitative models of sequence-function relationships [e.g., 44].

See Supplemental Information for full derivations of the mathematical results presented above. Python code implementing the embedding distillation algorithm described the section “Computational analysis of generalized one-hot models” and used for generating Fig. 4 is available at https://github.com/jbkinney/24_posfai2.

## ACKNOWLEDGMENTS

We thank Peter Koo and Vijay Balasubramanian for helpful discussions. This work was supported by NIH grant R35 GM133613 (AP, DMM), NIH grant R35 GM133777 (AP, JBK), NIH grant R01 HG011787 (JBK), the Alfred P. Sloan foundation (DMM), and additional funding from the Simons Center for Quantitative Biology at CSHL (DMM, JBK).

