## Supplemental Information for "Symmetry, gauge freedoms, and the interpretability of sequence-function Relationships"

March 17, 2025

### 1 Preliminaries

#### 1.1 Notation

- $V$ denotes feature space, i.e., the vector space in which embeddings and parameters live.
- $M$ denotes the dimension feature space.
- $\vec{x}$ denotes the sequence embedding of a model.
- $\vec{\theta}$ denotes the parameters of a model.
- $f$ denotes a linear model.
- $\gamma$ denotes the number of gauge freedoms.
- $G$ denotes the space of gauge freedoms.
- $\Theta$ denotes a gauge space.
- $H$ denotes a group.
- $\leftrightarrow$ denotes a transposition.
- $R$ denotes a group representation.
- $\simeq$ denotes equivalence.
- $\mathcal{U}$ denotes a finite set.
- $\alpha$ denotes the number of characters in an alphabet.
- $c$ denotes a character in an alphabet.
- $L$ denotes the length of a sequence.

**Note**: We assume throughout this document that $V$ is complex, whereas Main Text assumes that $V$ is real in order to simplify the presentation. Sec. 5.3 shows that our embedding distillation procedure, derived here assuming complex $V$, still works when $V$ is restricted to the reals.

#### 1.2 Specific sets

- $\mathcal{I}_\alpha$ denotes $(1, 2, \dots, \alpha)$, an ordered set comprising the first $\alpha$ positive integers.
- $\mathcal{A}$ denotes the alphabet $(c_1, \dots, c_\alpha)$, an ordered set comprising $\alpha$ characters. Note that $\mathcal{A}$ is written as an unordered set in main text.
- $\mathcal{S}$ denotes sequence space, i.e., the set of sequences of length $L$ built from the $\alpha$ characters in $\mathcal{A}$.

#### 1.3 Specific groups

- $S_\alpha$ denotes the symmetric group and acts on $\mathcal{I}_\alpha$.
- $H_{\text{CP}}$ denotes the group of character permutations and acts on $\mathcal{A}$.
- $H^l_{\text{CP}}$ (written $H_l$ in Main Text) denotes the group of single-position character permutations and acts on $\mathcal{S}$.
- $H_{\text{PSCP}}$ denotes the group of position-specific character permutations and acts on $\mathcal{S}$.
- $H_{\text{PP}}$ denotes the group of position permutations and acts on $\mathcal{S}$.
- $H_{\text{Ham}}$ denotes the group of Hamming graph symmetries and acts on $\mathcal{S}$.

#### 1.4 Specific embeddings and representations

- $\vec{x}^{\,\text{triv}}$ denotes the trivial embedding of $\mathcal{I}_\alpha$, $\mathcal{A}$, or $\mathcal{S}$. $R^{\,\text{triv}}$ denotes the trivial representation, under which $\vec{x}^{\,\text{triv}}$ transforms.
- $\vec{x}^{\,\text{ohe}}$ denotes the one-hot embedding of $\mathcal{I}_\alpha$ or $\mathcal{A}$. $R^{\,\text{ohe}}$ denotes the one-hot representation, under which $\vec{x}^{\,\text{ohe}}$ transforms.
- $\vec{x}^{\,\text{sim}}$ denotes the simplex embedding of $\mathcal{I}_\alpha$ or $\mathcal{A}$. $R^{\,\text{sim}}$ denotes the simplex representation, under which $\vec{x}^{\,\text{sim}}$ transforms.
- $\vec{x}^{\,\text{tet}}$ denotes the tetrahedral embedding of $\mathcal{A}$, applicable when $\alpha = 4$. $R^{\,\text{tet}}$ denotes the tetrahedral representation, under which $\vec{x}^{\,\text{tet}}$ transforms.
- $\vec{x}^{\text{ohe}}_l$, $l \in \mathcal{I}_L$, denote a single-position one-hot embedding of $\mathcal{S}$. $R^{\text{ohe}}_l$ denotes the representation under which $\vec{x}^{\text{ohe}}_l$ transforms.
- $\vec{x}^{\text{sim}}_l$, $l \in \mathcal{I}_L$, denotes a single-position simplex embedding of $\mathcal{S}$. $R^{\text{sim}}_l$ denotes the simplex representation, under which $\vec{x}^{\text{sim}}_l$ transforms.

#### 1.5 Definitions

- **co-transformation**: Two equivariant embeddings *co-transform* if they are transformed by the same group representation.
- **embedding**: An *embedding* $\vec{x}$ is a map from a set $\mathcal{U}$ to a vector space $\mathbb{R}^m$.
- **equivariance**:
  - An embedding $\vec{x}$ of a set $\mathcal{U}$ is *equivariant* under a group $H$ iff there is a group representation $R$ such that $\vec{x}(hu) = R(h)\vec{x}(u)$ for all $h \in H$ and all $u \in \mathcal{U}$.
  - A linear model $f$ on a set $\mathcal{U}$ is *equivariant* under a group $H$ iff $f(u, \vec{\theta}) = \vec{\theta}^{\,\dagger}\vec{x}(u)$ for all $u \in \mathcal{U}$, where $\vec{x}$ is an embedding of $\mathcal{U}$ that is equivariant under $H$, and $\vec{\theta} \in \mathbb{R}^m$ where $m = \dim \vec{x}$.
- **equivalence**:
  - Two embeddings $\vec{x}$ and $\vec{y}$ of a set $\mathcal{U}$ are *equivalent*, denoted $\vec{x} \simeq \vec{y}$, iff there is an invertible matrix $A$ (a similarity transformation) such that $\vec{y}(u) = A\vec{x}(u)$ for all $u \in \mathcal{U}$.
  - Two representations $R$ and $R'$ of a group $H$ are *equivalent*, denoted $R \simeq R'$, iff there is an invertible matrix $A$ (a similarity transformation) such that $R(h) = AR'(h)A^{-1}$ for all $h \in H$.
- **degree**: The degree of a representation $R$, denoted $\deg R$, is the number of rows (or equivalently the number of columns) of each matrix in the representation.
- **full rank embedding**: An embedding $\vec{x}$ of degree $m$ is *full rank* iff $\text{span}\, \vec{x} = \mathbb{R}^m$.
- **irreducibility**: An *irreducible* embedding is an equivariant embedding that transforms under an irreducible representation.
- **nonzero embedding**: An embedding $\vec{x}$ of a set $\mathcal{U}$ is *nonzero* iff $\vec{x}(u) \neq 0$ for some $u \in \mathcal{U}$.

- **linear model**: A complex-valued model of a sequence-function relationship is a function of the form $f(s\vec{\theta}) = \vec{\theta}^\dagger \vec{x}(s)$, where $\vec{x} : \mathcal{S} \to \mathbb{C}^M$ and $\vec{\theta} \in \mathbb{C}^M$.
- **module**:
  - A *module* is a vector space, together with a group representation that transforms the elements of that vector space.
  - The *module* of an equivariant embedding $\vec{x}$ that has degree $m$ and transforms under the representation $R$ is the module defined by the vector space $\mathbb{R}^m$ and representation $R$.
- **representation**: A group representation $R$ is a map from a group $H$ to a set of matrices that preserve the multiplicaiton rules of $H$, i.e., for which $R(h_1)R(h_2) = R(h_1 h_2)$ for all $h_1, h_2 \in H$.
- **span of an embedding**: The *span* of an embedding $\vec{x}$ of a set $\mathcal{U}$, denoted span $\vec{x}$, is the span of the set of vectors $\{\vec{x}(u) : u \in \mathcal{U}\}$.

### 2 Single-character embeddings and representations

In what follows, we consider a variety of singe-character embeddings and their representations. For the general definitions we assume an alphabet $\mathcal{A} = (c_1, \dots, c_\alpha)$; for the DNA-specific examples we assume an alphabet $\mathcal{A}_{\text{DNA}} = (\text{A}, \text{C}, \text{G}, \text{T})$.

#### 2.1 The trivial embedding $\vec{x}^{\text{triv}}$ and representation $R^{\text{triv}}$

The trivial embedding $\vec{x}^{\text{triv}}$ of characters in $\mathcal{A}$ has dimension 1 and is given by

$$\vec{x}^{\text{triv}}(c) = [1] \quad \text{for all} \quad c \in \mathcal{A}. \tag{1}$$

$\vec{x}^{\text{triv}}$ transforms under the trivial representation, $R^{\text{triv}}$, of $H_{\text{CP}}$. $R^{\text{triv}}$ has degree 1 and is given by

$$R^{\text{triv}}(h) = [1] \quad \text{for all} \quad h \in H_{\text{CP}}. \tag{2}$$

#### 2.2 The one-hot embedding $\vec{x}^{\text{ohe}}$ and representation $R^{\text{ohe}}$

The one-hot embedding, $\vec{x}^{\text{ohe}}$, is defined to be an $\alpha$-dimensional vector having elements

$$[\vec{x}^{\text{ohe}}(c)]_i = \begin{cases} 1 & \text{if } c_i = c, \\ 0 & \text{otherwise.} \end{cases} . \tag{3}$$

For example, $\vec{x}^{\text{ohe}}$ has dimension 4 for DNA alphabet $\mathcal{A}_{\text{DNA}}$ and is given by

$$\vec{x}^{\text{ohe}}(\text{A}) = \begin{bmatrix} 1 \\ 0 \\ 0 \\ 0 \end{bmatrix}, \quad \vec{x}^{\text{ohe}}(\text{C}) = \begin{bmatrix} 0 \\ 1 \\ 0 \\ 0 \end{bmatrix}, \quad \vec{x}^{\text{ohe}}(\text{G}) = \begin{bmatrix} 0 \\ 0 \\ 1 \\ 0 \end{bmatrix}, \quad \vec{x}^{\text{ohe}}(\text{T}) = \begin{bmatrix} 0 \\ 0 \\ 0 \\ 1 \end{bmatrix}. \tag{4}$$

$\vec{x}^{\text{ohe}}$ transforms under the one-hot representation, $R^{\text{ohe}}$, of $H_{\text{CP}}$. $R^{\text{ohe}}$ is isomorphic to what is called the "defining representation" of the symmetric group, but here we use the term "one-hot representation" for consistency. $H_{\text{CP}}$ is generated by transpositions, i.e., the exchange of two specific characters. We denote each transposition as $c \leftrightarrow c'$, where $c, c'$ are distinct characters in $\mathcal{A}$. For any transposition $c \leftrightarrow c'$, the corresponding one-hot representation has columns given by

$$[R^{\text{ohe}}(c \leftrightarrow c')]_{:,j} = \begin{cases} \vec{x}^{\text{ohe}}(c') & \text{if } c_j = c, \\ \vec{x}^{\text{ohe}}(c) & \text{if } c_j = c', \\ \vec{x}^{\text{ohe}}(c_j) & \text{otherwise.} \end{cases} . \tag{5}$$

In the case of $\mathcal{A}_{\text{DNA}}$, $H_{\text{CP}}$ is generated by 6 transpositions. For these transpositions, the one-hot representations are

$$R^{\text{ohe}}(\text{A}\leftrightarrow\text{C}) = \begin{bmatrix} 0 & 1 & 0 & 0 \\ 1 & 0 & 0 & 0 \\ 0 & 0 & 1 & 0 \\ 0 & 0 & 0 & 1 \end{bmatrix}, \quad R^{\text{ohe}}(\text{A}\leftrightarrow\text{G}) = \begin{bmatrix} 0 & 0 & 1 & 0 \\ 0 & 1 & 0 & 0 \\ 1 & 0 & 0 & 0 \\ 0 & 0 & 0 & 1 \end{bmatrix}, \quad R^{\text{ohe}}(\text{A}\leftrightarrow\text{T}) = \begin{bmatrix} 0 & 0 & 0 & 1 \\ 0 & 1 & 0 & 0 \\ 0 & 0 & 1 & 0 \\ 1 & 0 & 0 & 0 \end{bmatrix}, \tag{6}$$

$$R^{\text{ohe}}(\text{C}\leftrightarrow\text{G}) = \begin{bmatrix} 1 & 0 & 0 & 0 \\ 0 & 0 & 1 & 0 \\ 0 & 1 & 0 & 0 \\ 0 & 0 & 0 & 1 \end{bmatrix}, \quad R^{\text{ohe}}(\text{C}\leftrightarrow\text{T}) = \begin{bmatrix} 1 & 0 & 0 & 0 \\ 0 & 0 & 0 & 1 \\ 0 & 0 & 1 & 0 \\ 0 & 1 & 0 & 0 \end{bmatrix}, \quad R^{\text{ohe}}(\text{G}\leftrightarrow\text{T}) = \begin{bmatrix} 1 & 0 & 0 & 0 \\ 0 & 1 & 0 & 0 \\ 0 & 0 & 0 & 1 \\ 0 & 0 & 1 & 0 \end{bmatrix}. \tag{7}$$

#### 2.3 The simplex embedding $\vec{x}^{\,\mathrm{sim}}$ and representation $R^{\,\mathrm{sim}}$

The simplex embedding and representation can be formulated in multiple equivalent ways. Here and in Main Text we use one formulation that is particularly amenable to analytic calculations: we define the simplex embedding $\vec{x}^{\,\mathrm{sim}}$ of $\mathcal{A}$ to be an $(\alpha-1)$-dimensional vector having elements

$$[\vec{x}_l^{\,\mathrm{sim}}(s)]_{i-1} = \begin{cases} 1 & \text{if } s_l = c_i \text{ and } i \neq \alpha, \\ -1 & \text{if } s_l = c_\alpha, \\ 0 & \text{otherwise.} \end{cases} \tag{8}$$

In the case of $\mathcal{A}_{\mathrm{DNA}}$, $\vec{x}^{\,\mathrm{sim}}$ has dimension 3 and is given by

$$\vec{x}^{\,\mathrm{sim}}(\mathrm{A}) = \begin{bmatrix} 1 \\ 0 \\ 0 \end{bmatrix}, \quad \vec{x}^{\,\mathrm{sim}}(\mathrm{C}) = \begin{bmatrix} 0 \\ 1 \\ 0 \end{bmatrix}, \quad \vec{x}^{\,\mathrm{sim}}(\mathrm{G}) = \begin{bmatrix} 0 \\ 0 \\ 1 \end{bmatrix}, \quad \vec{x}^{\,\mathrm{sim}}(\mathrm{T}) = \begin{bmatrix} -1 \\ -1 \\ -1 \end{bmatrix}. \tag{9}$$

$\vec{x}^{\,\mathrm{sim}}$ transforms under the simplex representation, $R^{\,\mathrm{sim}}$, of $H_{\mathrm{CP}}$. $R^{\,\mathrm{sim}}$ is isomorphic to the "standard representation" of the symmetric group, but here we use the term "simplex representation" for consistency. For any transposition $c \leftrightarrow c'$, the corresponding simplex representation has columns given by

$$[R^{\,\mathrm{sim}}(c \leftrightarrow c')]_{:,j} = \begin{cases} \vec{x}^{\,\mathrm{sim}}(c') & \text{if } c_j = c, \\ \vec{x}^{\,\mathrm{sim}}(c) & \text{if } c_j = c', \\ \vec{x}^{\,\mathrm{sim}}(c_j) & \text{otherwise.} \end{cases} . \tag{10}$$

In the case of $\mathcal{A}_{\mathrm{DNA}}$, the representations of the 6 generating elements of $H_{\mathrm{CP}}$ are

$$R^{\,\mathrm{sim}}(\mathrm{A} \leftrightarrow \mathrm{C}) = \begin{bmatrix} 0 & 1 & 0 \\ 1 & 0 & 0 \\ 0 & 0 & 1 \end{bmatrix}, \qquad R^{\,\mathrm{sim}}(\mathrm{A} \leftrightarrow \mathrm{G}) = \begin{bmatrix} 0 & 0 & 1 \\ 0 & 1 & 0 \\ 1 & 0 & 0 \end{bmatrix}, \qquad R^{\,\mathrm{sim}}(\mathrm{C} \leftrightarrow \mathrm{G}) = \begin{bmatrix} 1 & 0 & 0 \\ 0 & 0 & 1 \\ 0 & 1 & 0 \end{bmatrix}, \tag{11}$$

$$R^{\,\mathrm{sim}}(\mathrm{A} \leftrightarrow \mathrm{T}) = \begin{bmatrix} -1 & 0 & 0 \\ -1 & 1 & 0 \\ -1 & 0 & 1 \end{bmatrix}, \qquad R^{\,\mathrm{sim}}(\mathrm{C} \leftrightarrow \mathrm{T}) = \begin{bmatrix} 1 & -1 & 0 \\ 0 & -1 & 0 \\ 0 & -1 & 1 \end{bmatrix}, \qquad R^{\,\mathrm{sim}}(\mathrm{G} \leftrightarrow \mathrm{T}) = \begin{bmatrix} 1 & 0 & -1 \\ 0 & 1 & -1 \\ 0 & 0 & -1 \end{bmatrix}. \tag{12}$$

#### 2.4 The tetrahedral embedding $\vec{x}^{\,\mathrm{tet}}$ and representation $R^{\,\mathrm{tet}}$

It is worth noting that, instead of the simplex embedding for DNA characters, some studies (e.g., [3]) have proposed what we call the "tetrahedral embedding", $\vec{x}^{\,\mathrm{tet}}$. The tetrahedral embedding is given by

$$\vec{x}^{\,\mathrm{tet}}(\mathrm{A}) = \begin{bmatrix} 1 \\ 1 \\ 1 \end{bmatrix}, \quad \vec{x}^{\,\mathrm{tet}}(\mathrm{C}) = \begin{bmatrix} 1 \\ -1 \\ -1 \end{bmatrix}, \quad \vec{x}^{\,\mathrm{tet}}(\mathrm{G}) = \begin{bmatrix} -1 \\ 1 \\ -1 \end{bmatrix}, \quad \vec{x}^{\,\mathrm{tet}}(\mathrm{T}) = \begin{bmatrix} -1 \\ -1 \\ 1 \end{bmatrix}. \tag{13}$$

$\vec{x}^{\,\mathrm{tet}}$ transforms under the tetrahedral representation, $R^{\,\mathrm{tet}}$, of $H_{\mathrm{CP}}$. In the tetrahedral representation, the 6 generating elements of $H_{\mathrm{CP}}$ are

$$R^{\,\mathrm{tet}}(\mathrm{A} \leftrightarrow \mathrm{C}) = \begin{bmatrix} 1 & 0 & 0 \\ 0 & 0 & -1 \\ 0 & -1 & 0 \end{bmatrix}, \qquad R^{\,\mathrm{tet}}(\mathrm{A} \leftrightarrow \mathrm{G}) = \begin{bmatrix} 0 & 0 & -1 \\ 0 & 1 & 0 \\ -1 & 0 & 0 \end{bmatrix}, \qquad R^{\,\mathrm{tet}}(\mathrm{A} \leftrightarrow \mathrm{T}) = \begin{bmatrix} 0 & -1 & 0 \\ -1 & 0 & 0 \\ 0 & 0 & 1 \end{bmatrix}, \tag{14}$$

$$R^{\,\mathrm{tet}}(\mathrm{C} \leftrightarrow \mathrm{G}) = \begin{bmatrix} 0 & 1 & 0 \\ 1 & 0 & 0 \\ 0 & 0 & 1 \end{bmatrix}, \qquad R^{\,\mathrm{tet}}(\mathrm{C} \leftrightarrow \mathrm{T}) = \begin{bmatrix} 0 & 0 & 1 \\ 0 & 1 & 0 \\ 1 & 0 & 0 \end{bmatrix}, \qquad R^{\,\mathrm{tet}}(\mathrm{G} \leftrightarrow \mathrm{T}) = \begin{bmatrix} 1 & 0 & 0 \\ 0 & 0 & 1 \\ 0 & 1 & 0 \end{bmatrix}. \tag{15}$$

The tetrahedral embedding is equivalent to the simplex embedding, and the tetrahedral representation is equivalent to the simplex representation:

$$A\vec{x}^{\,\mathrm{sim}} = \vec{x}^{\,\mathrm{tet}} \quad \text{and} \quad AR^{\,\mathrm{sim}}A^{-1} = R^{\,\mathrm{tet}} \quad \text{where} \quad A = \begin{bmatrix} 1 & 1 & -1 \\ 1 & -1 & 1 \\ 1 & -1 & -1 \end{bmatrix}. \tag{16}$$

#### 2.5 Maschke decompositions of $\vec{x}^{\,\text{ohe}}$ and $R^{\text{ohe}}$.

The one-hot representation decomposes, by Maschke's Theorem, into the direct sum of the trivial representation and the simplex representation:

$$R^{\text{ohe}} \simeq R^{\text{triv}} \oplus R^{\text{sim}}. \tag{17}$$

Similarly, the one-hot embedding decomposes into the direct sum of the trivial embedding and the simplex embedding:

$$\vec{x}^{\,\text{ohe}} \simeq \vec{x}^{\,\text{triv}} \oplus \vec{x}^{\,\text{sim}}. \tag{18}$$

In the case of $\mathcal{A}_{\text{DNA}}$, these decompositions are accomplished by the similarity transformation matrix $T$, in the sense that

$$T\,R^{\text{ohe}}\,T^{-1} = \left[\begin{array}{c|ccc} R^{\text{triv}} & 0 & 0 & 0 \\ \hline 0 & & & \\ 0 & & R^{\text{sim}} & \\ 0 & & & \end{array}\right], \quad T\,\vec{x}^{\,\text{ohe}} = \left[\begin{array}{c} \vec{x}^{\,\text{triv}} \\ \hline \\ \vec{x}^{\,\text{sim}} \\ \\ \end{array}\right], \tag{19}$$

where

$$T = \begin{bmatrix} 1 & 1 & 1 & 1 \\ 1 & 0 & 0 & -1 \\ 0 & 1 & 0 & -1 \\ 0 & 0 & 1 & -1 \end{bmatrix} \quad \Rightarrow \quad T^{-1} = \frac{1}{4}\begin{bmatrix} 1 & 3 & -1 & -1 \\ 1 & -1 & 3 & -1 \\ 1 & -1 & -1 & 3 \\ 1 & -1 & -1 & -1 \end{bmatrix}. \tag{20}$$

The Maschke decomposition formulas for $R^{\text{ohe}}$ and $\vec{x}^{\,\text{ohe}}$, using $R^{\text{tet}}$ and $\vec{x}^{\,\text{tet}}$ in place of $R^{\text{sim}}$ and $\vec{x}^{\,\text{sim}}$, are given in terms of the alternative decomposition matrix $T_{\text{alt}}$ by,

$$T_{\text{alt}}\,R^{\text{ohe}}\,T_{\text{alt}}^{-1} = \left[\begin{array}{c|ccc} R^{\text{triv}} & 0 & 0 & 0 \\ \hline 0 & & & \\ 0 & & R^{\text{tet}} & \\ 0 & & & \end{array}\right] \quad \text{and} \quad T_{\text{alt}}\,\vec{x}^{\,\text{ohe}} = \left[\begin{array}{c} \vec{x}^{\,\text{triv}} \\ \hline \\ \vec{x}^{\,\text{tet}} \\ \\ \end{array}\right], \tag{21}$$

where

$$T_{\text{alt}} = \begin{bmatrix} 1 & 1 & 1 & 1 \\ 1 & 1 & -1 & -1 \\ 1 & -1 & 1 & -1 \\ 1 & -1 & -1 & 1 \end{bmatrix} \quad \Rightarrow \quad T_{\text{alt}}^{-1} = \frac{1}{4}\begin{bmatrix} 1 & 1 & 1 & 1 \\ 1 & 1 & -1 & -1 \\ 1 & -1 & 1 & -1 \\ 1 & -1 & -1 & 1 \end{bmatrix}. \tag{22}$$

Note that $T_{\text{alt}}$ is a Haddamard matrix of order 4 [3]. Also note that, unlike the simplex embedding, it is unclear how to generalize the tetrahedral embedding to alphabets that contain arbitrary numbers of characters.

### 3 Results for general groups

#### 3.1 Nonzero embeddings

**Claim 3.1.** *Let $\vec{x}$ be an embedding of a set $\mathcal{U}$ that transforms under a representation of a group $H$ that is transitive on $\mathcal{U}$. Then either $\vec{x}(u) = \vec{0}$ all $u \in \mathcal{U}$, or $\vec{x}(u) \neq \vec{0}$ for all $u \in \mathcal{U}$.*

*Proof.* Assume that $\vec{x}(u) = \vec{0}$ for some $u \in \mathcal{U}$, and choose any other $u' \in \mathcal{U}$. Because $H$ is transitive, there is a group element $h \in \mathcal{H}$ such that $u' = hu$. Letting $R$ denote the representation of $H$ that $\vec{x}$ transforms under, we find

$$\vec{x}(u') = R(h)\vec{x}(u) = R(h)\vec{0} = \vec{0}. \tag{23}$$

We thus find that $\vec{x}(u) = \vec{0}$ for some $u \in \mathcal{U}$ implies that $\vec{x}(u) = \vec{0}$ for all $u \in \mathcal{U}$. This completes the proof. We note, however, that if $H$ is not transitive on $\mathcal{U}$, then Claim 3.1 will generally not hold. □

#### 3.2 Equivariance

**Claim 3.2.** *If $f$ is a linear model based on the embedding $\vec{x}$ of a set $\mathcal{U}$ into a vector space $V$, and $\vec{x}$ is an equivariant embedding that transforms under a representation $R$ of a group $H$, then*

$$f\left(u; \vec{\theta}\,\right) = f\left(hu; R(h)^{-1\dagger}\,\vec{\theta}\,\right) \tag{24}$$

*for all $\vec{\theta} \in V$, all $u \in \mathcal{U}$, and all $h \in H$.*

*Proof.*

$$
\begin{aligned}
f(hu; R(h)^{-1\dagger}\vec{\theta}) &= [R(h)^{-1\dagger}\vec{\theta}]^\dagger \vec{x}(hu) && (25)\\
&= \vec{\theta}^\dagger R(h)^{-1} R(h)\vec{x}(u) && (26)\\
&= \vec{\theta}^\dagger \vec{x}(u) && (27)\\
&= f(u; \vec{\theta}). && (28)
\end{aligned}
$$

□

#### 3.3 Irreducible embeddings are full rank

**Claim 3.3.** *Every nonzero equivariant embedding that transforms under an irreducible representation is full rank.*

*Proof.* Let $\vec{x}$ be a nonzero embedding of a set $\mathcal{U}$ into a vector space $V$. Further assume that $\vec{x}$ transforms under an irreducible representation $R$ of a group $H$. Since $R$ transforms vectors in $V$, $V$ is an irreducible $H$ module. For any $h \in H$,

$$
\begin{aligned}
R(h)\,\mathrm{span}\,\vec{x} &= \mathrm{span}\,\{R(h)\vec{x}(u) : u \in \mathcal{U}\} && (29)\\
&= \mathrm{span}\,\{\vec{x}(hu) : u \in \mathcal{U}\} && (30)\\
&= \mathrm{span}\,\{\vec{x}(u) : u \in \mathcal{U}\} && (31)\\
&= \mathrm{span}\,\vec{x}. && (32)
\end{aligned}
$$

Therefore, $\mathrm{span}\,\vec{x}$ is an $H$-invariant subspace of $V$. Since $V$ is an irreducible module, the only $H$-invariant subspaces of $V$ are 0 and $V$. Because $\vec{x}$ is nonzero, $\mathrm{span}\,\vec{x} \neq 0$. We thus find that $\mathrm{span}\,\vec{x} = V$, proving that $\vec{x}$ is full rank. □

#### 3.4 Direct sums of inequivalent irreducible embeddings have full rank (Theorem 2 of Main Text)

**Claim 3.4.** *(Theorem 2 of Main Text) For each $k \in \{1, \ldots, K\}$, let $\vec{x}_k$ be a nonzero embedding that transforms under an irreducible representation $R_k$ of a group $H$. Then the direct sum of all $\vec{x}_k$ is full rank if all $R_k$ are pairwise inequivalent.*

*Proof.* Let $V = \bigoplus_{k=1}^K V_k$ where $V_k$ is the vector space to which the embedding $\vec{x}_k$ maps. Letting $R = \bigoplus_{k=1}^K R_k$, we see that $R$ is a representation of $H$ over $V$. Given $R$, define an $H$-invariant inner product $\langle \cdot, \cdot \rangle$ over $V$, i.e. an inner product that satisfies

$$\langle R(h)\vec{v}, R(h)\vec{w}\rangle = \langle \vec{v}, \vec{w}\rangle \quad \text{for all} \quad \vec{v}, \vec{w} \in V \quad \text{and} \quad h \in H. \tag{33}$$

Such an inner product can always be constructed (e.g., see p. 16 of [2]). Next, using the notation $\vec{x}(s) = \bigoplus_{k=1}^K \vec{x}_k(s)$, define the subspace $W$ of $V$ given by

$$W = \{\vec{w} \in V : \langle \vec{w}, \vec{x}(s)\rangle = 0 \quad \text{for all} \quad s \in \mathcal{S}\}. \tag{34}$$

$W$ is $H$-invariant, since for all $\vec{w} \in W$ and $h \in H$,

$$\langle R(h)\vec{w}, \vec{x}(s)\rangle = \left\langle \vec{w}, R(h)^{-1}\vec{x}(s)\right\rangle = \left\langle \vec{w}, \vec{x}(h^{-1}s)\right\rangle = 0. \tag{35}$$

We are goingto show that $\dim(W) = 0$.

We start by showing that the projection operator $P$ that projects $V$ orthogonally on to $W$ according to the inner product $\langle \cdot, \cdot \rangle$ is an $H$-homomorphism. Assume $W$ has dimension $n$, and let $\{\hat{e}_1, \ldots, \hat{e}_n\}$ be an orthonormal basis of $W$ (i.e., $\langle \hat{e}_i, \hat{e}_j\rangle = \delta_{ij}$ for all $i, j \in \mathcal{I}_n$). We define the projection operator $P$ to be such that

$$P\vec{v} = \sum_{i=1}^n \hat{e}_i \langle \hat{e}_i, \vec{v}\rangle \quad \text{for all} \quad v \in V. \tag{36}$$

Because $\langle \cdot, \cdot \rangle$ is a $H$-invariant inner product, $\langle R(h)\hat{e}_i, R(h)\hat{e}_j\rangle = \delta_{ij}$ for all $h \in H$, and thus $\{R(h)\hat{e}_i\}_{i=1}^n$ is also an orthonormal basis for $W$. Consequently, for all $h \in H$ and $\vec{v} \in V$,

$$P\vec{v} = \sum_{i=1}^n R(h)\hat{e}_i \langle R(h)\hat{e}_i, \vec{v}\rangle = R(h)\sum_{i=1}^n \hat{e}_i \left\langle \hat{e}_i, R(h)^{-1}\vec{v}\right\rangle = R(h)PR(h)^{-1}\vec{v}. \tag{37}$$

We therefore see that $P$ commutes with $R$. $P$ is therefore an $H$-homomorphism.

By theorem 1.7.8 in [2], there is a set of real scalars $\{c_k\}_{k=1}^K$ such that

$$P = \bigoplus_{k=1}^{K} c_k I_k, \tag{38}$$

where $I_k$ denotes the identity operator on $V_k$. Because $P\vec{x}(s) = \vec{0}$ for all $s \in \mathcal{S}$,

$$0 = \langle P\vec{x}(s), P\vec{x}(s)\rangle = \sum_{k=1}^{K} \langle c_k\vec{x}_k(s), c_k\vec{x}_k(s)\rangle = \sum_{k=1}^{K} ||c_k||^2 ||\vec{x}_k(s)||^2. \tag{39}$$

Since all $\vec{x}_k$ are nonzero, this is only possible if all $c_k = 0$. We thus find that $P$ projects all vectors to $\vec{0}$, and hence that $W$ has dimension 0. Since $W$ is the orthogonal complement of span $\vec{x}$, $\vec{x}$ has full rank. This completes the proof. □

### 4 Results for $S_\alpha$, the symmetric group

The symmetric group, $S_\alpha$, is the group of permutations among the integers $\mathcal{I}_\alpha$. $S_\alpha$ is generated by transpositions, which exchange two elements of $\mathcal{I}_\alpha$ while leaving all other elements unchanged. In what follows, we use $i \leftrightarrow j$ to denote the transposition that exchanges elements $i, j \in \mathcal{I}_\alpha$.

#### 4.1 The trivial embedding $\vec{x}^{\,\text{triv}}$ and representation $R^{\,\text{triv}}$

The trivial embedding of $\mathcal{I}_\alpha$, $\vec{x}^{\,\text{triv}}$, has degree 1 and is given by

$$\vec{x}^{\,\text{triv}}(i) = [1] \ \text{ for all } \ i \in I_\alpha. \tag{40}$$

$\vec{x}^{\,\text{triv}}$ transforms under the trivial representation, $R^{\,\text{triv}}$, which also has degree 1 and is given by

$$R^{\,\text{triv}}(h) = [1] \ \text{ for all } \ h \in S_\alpha. \tag{41}$$

#### 4.2 The simplex embedding $\vec{x}^{\,\text{sim}}$ and representation $R^{\,\text{sim}}$

Here we define an explicit coordinate realization of the simplex embedding $\vec{x}^{\,\text{sim}}$ of $\mathcal{I}_\alpha$, as well as the simplex representation $R^{\,\text{sim}}$ of $S_\alpha$, under which $\vec{x}^{\,\text{sim}}$ transforms. Both $\vec{x}^{\,\text{sim}}$ and $R^{\,\text{sim}}$ have degree $\alpha - 1$, and we use $k, l \in \mathcal{I}_{\alpha-1}$ to index the elements of $\vec{x}^{\,\text{sim}}$ and $R^{\,\text{sim}}$. We define the coordinates of $\vec{x}^{\,\text{sim}}$ to be

$$[\vec{x}^{\,\text{sim}}(i)]_k = \delta_{ik} \ \text{ for } \ i \in \mathcal{I}_{\alpha-1}, \tag{42}$$

$$[\vec{x}^{\,\text{sim}}(\alpha)]_k = -1. \tag{43}$$

Correspondingly, we define the coordinate realization of $R^{\,\text{sim}}$ on transpositions to be

$$[R^{\,\text{sim}}(i \leftrightarrow j)]_{kl} = \delta_{kl} - \delta_{ik}\delta_{il} - \delta_{jk}\delta_{jl} + \delta_{ik}\delta_{jl} + \delta_{jk}\delta_{il}, \tag{44}$$

$$[R^{\,\text{sim}}(i \leftrightarrow \alpha)]_{kl} = \delta_{kl} - \delta_{kl}\delta_{ik} - \delta_{il}. \tag{45}$$

One can readily verify that these definitions work by showing that $\vec{x}^{\,\text{sim}}$ does in fact transform according to $R^{\,\text{sim}}$ for all transpositions in $S_\alpha$, and thereby showing that $R^{\,\text{sim}}$ is indeed a valid representation of $S_\alpha$. The simplex embedding and representation in Sec. 2.3 is an example of this formulation for $\alpha = 4$. Finally, we note that $\vec{x}^{\,\text{sim}}$ has the useful property that

$$\sum_{i=1}^{\alpha} \vec{x}^{\,\text{sim}}(i) = \vec{0}. \tag{46}$$

#### 4.3 $\vec{x}^{\,\text{triv}}$ and $\vec{x}^{\,\text{sim}}$ are the only inequivalent irreducible embeddings that transform under $S_\alpha$

**Claim 4.1.** *If a nonzero embedding of $\mathcal{I}_\alpha$ transforms under an irreducible representation of $S_\alpha$, then it is equivalent to either the trivial embedding or the simplex embedding.*

*Proof.* Assume that $\vec{x}$ is a nonzero embedding of $\mathcal{I}_\alpha$, that $\vec{x}$ has degree $m$, and that $\vec{x}$ transforms under an irreducible representation $R$ of $\mathcal{S}_\alpha$.

First consider $\alpha = 2$. $S_2$ has only two inequivalent irreducible representations: the trivial representation and the simplex representation. The trivial representation supports the embedding $\vec{x}^{\,\mathrm{triv}}$, and the simplex representation supports the simplex embedding $\vec{x}^{\,\mathrm{sim}}$. This proves the claim for $\alpha = 2$.

Now consider $\alpha \geq 3$. Select $i, j, k \in \mathcal{I}_\alpha$, where $i \neq j$, $i \neq k$, $j \neq k$, and assume that $\vec{x}(i) = \vec{x}(j) \neq \vec{x}(k)$. The transposition $j \leftrightarrow k$ exchanges $j$ and $k$ without affecting $i$. Therefore, the representation of $R(j \leftrightarrow k)$ must exchange the vectors $\vec{x}(j)$ and $\vec{x}(k)$ while leaving the vector $\vec{x}(i)$ invariant. But this is impossible because $\vec{x}(i) = \vec{x}(j)$. Consequently, $\vec{x}(i)$ must be either the same for all $i \in \mathcal{I}_\alpha$, or must be different for all $i \in \mathcal{I}_\alpha$. If $\vec{x}(i)$ is the same for all $i$, then $\vec{x} \simeq \vec{x}^{\,\mathrm{triv}}$. In the rest of this proof we assume that $\vec{x}(i)$ is different for all $i \in \mathcal{I}_\alpha$. We will show that this implies $\vec{x} \simeq \vec{x}^{\,\mathrm{sim}}$.

Claim 3.3 shows that, because $\vec{x}$ is irreducible, $\vec{x}$ is also full rank. We can therefore choose a subset $\{n_1, \ldots, n_m\} \subseteq \mathcal{I}_\alpha$ such that $\{\vec{x}(n_1), \ldots, \vec{x}(n_m)\}$ is a basis for $\mathbb{R}^m$. For simplicity, we assume that $n_i = i$ for all $i \in \mathcal{I}_m$, so that $\mathcal{B} = \{\vec{x}(i) : i \in \mathcal{I}_m\}$ is the basis; we revisit this assumption at the end of the proof. In the basis $\mathcal{B}$, the elements of $\vec{x}$ are

$$[\vec{x}(i)]_k = \delta_{ik} \tag{47}$$

for all $i, k \in \mathcal{I}_m$.

Now choose $a, b, j \in \mathcal{I}_m$ such that $a \neq b$, $a \neq j$, $b \neq j$. We can compute the elements of $R(a \leftrightarrow b)$ in basis $\mathcal{B}$ by noting that

$$[\vec{x}(a)]_i = \sum_{k=1}^{m} [R(a \leftrightarrow b)]_{ik} [\vec{x}(b)]_k = [R(a \leftrightarrow b)]_{ib}, \tag{48}$$

$$[\vec{x}(b)]_i = \sum_{k=1}^{m} [R(a \leftrightarrow b)]_{ik} [\vec{x}(a)]_k = [R(a \leftrightarrow b)]_{ia}, \tag{49}$$

$$[\vec{x}(j)]_i = \sum_{k=1}^{m} [R(a \leftrightarrow b)]_{ik} [\vec{x}(j)]_k = [R(a \leftrightarrow b)]_{ij} \quad \text{for} \quad j \neq a, b. \tag{50}$$

and therefore that,

$$[R(a \leftrightarrow b)]_{ia} = \delta_{ib}, \tag{51}$$

$$[R(a \leftrightarrow b)]_{ib} = \delta_{ia}, \tag{52}$$

$$[R(a \leftrightarrow b)]_{ij} = \delta_{ij} \quad \text{for} \quad j \neq a, b. \tag{53}$$

Combining Eq. 51, Eq. 52, and Eq. 53 into a single expression gives that, for any $a, b \in I_m$ such that $a \neq b$, and any $i, j \in I_m$,

$$[R(a \leftrightarrow b)]_{ij} = \delta_{ij}(1 - \delta_{aj})(1 - \delta_{bj}) + \delta_{aj}\delta_{bi} + \delta_{bj}\delta_{ai}, \tag{54}$$

$$= \delta_{ij} - \delta_{ai}\delta_{aj} - \delta_{bi}\delta_{bj} + \delta_{ai}\delta_{bj} + \delta_{bi}\delta_{aj}. \tag{55}$$

Now choose $c \in \mathcal{I}_\alpha \setminus \mathcal{I}_m$ and $a, j \in \mathcal{I}_m$ such that $a \neq j$. Because $\mathcal{B}$ is a basis, there are coefficients $r_i^c$, $i \in \mathcal{I}_m$, such that

$$\vec{x}(c) = \sum_{i=1}^{m} r_i^c \vec{x}(i) \tag{56}$$

Consequently, the elements of $\vec{x}(c)$ in the basis $\mathcal{B}$ are,

$$[\vec{x}(c)]_i = r_i^c. \tag{57}$$

We compute the matrix elements of $R(a \leftrightarrow c)$ by noting that

$$[\vec{x}(a)]_i = \sum_{k=1}^{m} [R(a \leftrightarrow c)]_{ik} [\vec{x}(c)]_k = \sum_{k=1}^{m} [R(a \leftrightarrow c)]_{ik} r_k^c, \tag{58}$$

$$[\vec{x}(c)]_i = \sum_{k=1}^{m} [R(a \leftrightarrow c)]_{ik} [\vec{x}(a)]_k = [R(a \leftrightarrow c)]_{ia}, \tag{59}$$

$$[\vec{x}(j)]_i = \sum_{k=1}^{m} [R(a \leftrightarrow c)]_{ik} [\vec{x}(j)]_k = [R(a \leftrightarrow c)]_{ij}. \tag{60}$$

From Eq. 47, Eq. 57, Eq. 59, and Eq. 60 we see that, for all $i, k \in \mathcal{I}_m$,

$$[R(a \leftrightarrow c)]_{ik} = \delta_{ik}(1 - \delta_{ak}) + r_i^c \delta_{ak} \tag{61}$$

are the elements of $R(a \leftrightarrow c)$. Plugging this into Eq. 58 then gives a constraint on the $r_i^c$ coefficients:

$$\delta_{ia} = \sum_{k=1}^{m} [\delta_{ik}(1 - \delta_{ak}) + r_i^c \delta_{ak}] r_k^c = r_i^c(1 - \delta_{ai}) + r_a^c r_i^c. \tag{62}$$

Considering different values for $i \in \mathcal{I}_\alpha$ in Eq. 62 then gives two specific constraints on $r_i^c$:

$$(r_a^c)^2 = 1 \text{ for } i = a \quad \text{and} \quad r_i^c(1 + r_a^c) = 0 \text{ for } i \neq a. \tag{63}$$

The left side of Eq. 63 gives $r_i^c = \pm 1$. If $r_a^c = 1$, the right side of Eq. 63 gives $r_i^c = 0$ for $i \neq a$. But this would mean that $\vec{x}(c) = \vec{x}(a)$, which violates the requirement that all $\vec{x}(i)$ are distinct. We thus find that $r_i^c = -1$. Therefore, for all $i, k \in \mathcal{I}_m$,

$$[\vec{x}(c)]_i = -1, \quad \text{and} \quad [R(a \leftrightarrow c)]_{ik} = \delta_{ik}(1 - \delta_{ak}) - \delta_{ak}. \tag{64}$$

In particular, this means that there is only a single possible $\vec{x}(c)$ for all $c \in \mathcal{I}_\alpha \setminus \mathcal{I}_m$. Since all $\vec{x}(c)$ must be distinct, there can only be one possible value for $c \geq m$, i.e. $c = \alpha = m + 1$. We therefore find that for $i, j, k, l \in I_{\alpha-1}$, $i \neq j$,

$$\begin{aligned}
[\vec{x}(i)]_k &= \delta_{ik} && (65)\\
[\vec{x}(\alpha)]_k &= -1 && (66)\\
[R(i \leftrightarrow j)]_{kl} &= \delta_{kl} - \delta_{ik}\delta_{il} - \delta_{jk}\delta_{jl} + \delta_{ik}\delta_{jl} + \delta_{jk}\delta_{il}. && (67)\\
[R(i \leftrightarrow \alpha)]_{kl} &= \delta_{kl} - \delta_{ik}\delta_{kl} - \delta_{il}. && (68)
\end{aligned}$$

We therefore see that $\vec{x} = \vec{x}^{\,\text{sim}}$ and $R = R^{\text{sim}}$. Finally, we revisit our simplifying assumption that $n_i = i$ for all $i \in \mathcal{I}_m$. It is readily seen that removing this assumption leads to

$$\vec{x} = T\vec{x}^{\,\text{sim}} \quad \text{and} \quad R = TR^{\text{sim}}T^{-1} \tag{69}$$

where $T$ is the similarity transformation that maps each $\vec{x}(i)$ to $\vec{x}(n_i)$, i.e.,

$$T = R(g) \quad \text{where} \quad g = \begin{pmatrix} 1 & 2 & \cdots & m \\ n_1 & n_2 & \cdots & n_m \end{pmatrix}. \tag{70}$$

Consequently, $\vec{x} \simeq \vec{x}^{\,\text{sim}}$ and $R \simeq R^{\text{sim}}$. This completes the proof. □

#### 4.4 Irreducible embeddings that co-transform under $S_\alpha$ are proportional

**Claim 4.2.** *Nonzero embeddings that transform under the same irreducible representation of the symmetric group are equal up to a constant of proportionality.*

*Proof.* Let $\vec{x}$ and $\vec{y}$ be two embeddings of $\mathcal{I}_\alpha$ that transform according to the same irreducible representation $R$ of $S_\alpha$. First we construct a matrix $A$ such that

$$\vec{y}(i) = A\vec{x}(i) \quad \text{for all} \quad i \in \mathcal{I}_\alpha. \tag{71}$$

From Claim 4.1, $\vec{x}$ and $\vec{y}$ must transform under the trivial representation or a representation equivalent to the simplex representation. If $\vec{x}$ and $\vec{y}$ transform under the trivial representation, then both $\vec{x}(i)$ and $\vec{y}(i)$ are nonzero scalars that do not depend on $i$, and Eq. 71 is satisfied by setting $A = \vec{y}/\vec{x}$. If $\vec{x}$ and $\vec{y}$ transform under a representation equivalent to the simplex representation, then both $\vec{x}$ and $\vec{y}$ have degree $m = \alpha - 1$. Define the $m \times m$ matrices $X = [\vec{x}(1)\ \vec{x}(2)\ \ldots\ \vec{x}(m)]$ and $Y = [\vec{y}(1)\ \vec{y}(2)\ \ldots\ \vec{y}(m)]$. Next define $A = YX^{-1}$; note that $X^{-1}$ exists because the columns of $X$ are linearly independent. By construction, $\vec{y}(i) = A\vec{x}(i)$ for $i \in I_{\alpha-1}$. Moreover $\sum_{i=1}^{\alpha} \vec{x}(i) = \sum_{i=1}^{\alpha} \vec{y}(i) = \vec{0}$ from Eq. 46. Therefore,

$$\vec{y}(\alpha) = -\sum_{i=1}^{\alpha-1} \vec{y}(i) = -\sum_{i=1}^{\alpha-1} A\vec{x}(i) = A\vec{x}(\alpha) \tag{72}$$

as well. This proves that Eq. 71 is satisfied when $\vec{x}$ and $\vec{y}$ transform under the simplex representation.

Next we show that $A$ commutes with $R$. For all $h \in \mathcal{S}_\alpha$ and all $i \in \mathcal{I}_\alpha$,

$$AR(h)\vec{x}(i) = A\vec{x}(hi) = \vec{y}(hi) = R(h)\vec{y}(i) = R(h)A\vec{x}(i). \tag{73}$$

Because $\vec{x}$ has full rank, this requires

$$AR(h) = R(h)A \tag{74}$$

for all $h \in S_\alpha$, thereby showing that $A$ and $R$ commute.

Finally, we show that $A = cI$ for some $c \in \mathbb{R}$. This is a consequence of Shur's Lemma (e.g., see Corollary 1.6.6 of ref. [2]), which tells us that any matrix that commutes with an irreducible representation is either invertible or the zero matrix. Now define $B = A - cI$ where $c$ is an eigenvalue of $A$. $B$ commutes with $R$, since

$$BR(h) = (A - cI)R(h) = R(h)(A - cI) = R(h)B. \tag{75}$$

But $B$ cannot be invertible because at least one of its eigenvalues is zero. $B$ must therefore be the zero matrix, implying that $A = cI$. This shows that $\vec{y} = c\vec{x}$, completing the proof. □

### 5 Results for $H_{\text{PSCP}}$, the group of position-specific character permutations

#### 5.1 Irreducible embeddings that co-transform under $H_{\text{PSCP}}$ are proportional (Theorem 1 of Main Text)

**Claim 5.1.** *(Theorem 1 of Main Text) Nonzero sequence embeddings that transform under the same irreducible representation of $H_{\text{PSCP}}$ are equal up to a constant of proportionality.*

*Proof.* As stated in Main Text, the group of position-specific character permutations, $H_{\text{PSCP}}$, is given by the direct product

$$H_{\text{PSCP}} = \bigotimes_{l=1}^{L} H_{\text{CP}}^{l}, \tag{76}$$

where $H_{\text{CP}}^{l}$ is the group of permutations among the $\alpha$ possible characters at position $l$ in a sequence. Therefore, every element $h \in H_{\text{PSCP}}$ can be expressed as the direct product

$$h = \bigotimes_{l=1}^{L} h_l, \quad h_l \in H_{\text{CP}}^{l}. \tag{77}$$

It is known that if we consider representations over an algebraically closed field like $\mathbb{C}$, then all irreducible representations of a product group can be expressed as tensor products of the irreducible representations of the individual group factors (e.g., Theorem 1.11.3 of ref. [2]). Therefore, any irreducible representation $R$ of $H_{\text{PSCP}}$ can be expressed as

$$R(h) = \bigotimes_{l=1}^{L} R_l(h_l), \tag{78}$$

where each $R_l$ is an irreducible representation of $H_{\text{CP}}^{l}$, and where, for each $h \in H_{\text{PSCP}}$, $h_l \in H_{\text{CP}}^{l}$ is the corresponding factor in Eq. 77.

Now let $\{\vec{x}_q\}_{q=1}^{Q}$ denote a set of embeddings that transform under $R$. Eq. 78 implies that each $\vec{x}_q$ must be expressible as a tensor product of site-specific embeddings as

$$\vec{x}_q = \bigotimes_{l=1}^{L} \vec{x}_q^{\,l}, \tag{79}$$

where each $\vec{x}_q^{\,l}$ is a site-specific embedding that transforms under $H_{\text{CP}}^{l}$. Because $H_{\text{CP}}^{l}$ is isomorphic to $S_\alpha$, each $R_l$ can be thought of as a representation of $\mathcal{S}_\alpha$, and each $\vec{x}_q^{\,l}$ can be thought of as an embedding of $\mathcal{I}_\alpha$. Claim 4.2 says that any two nonzero embeddings of $\mathcal{I}_\alpha$ that transform under the same irreducible representation $S_\alpha$ must be scalar multiples of one another. There must therefore exist, for every $l \in \mathcal{I}_L$, a nonzero embedding $\vec{x}^{\,l}$ that transforms under $R_l$ and that satisfies

$$\vec{x}_q^{\,l} = c_q^l \vec{x}^{\,l} \tag{80}$$

for all $q \in \mathcal{I}_Q$. We therefore find that

$$\vec{x}_q = c_q \vec{x}, \quad \text{where} \quad c_q = \prod_{l=1}^{L} c_q^l \quad \text{and} \quad \vec{x} = \bigotimes_{l=1}^{L} \vec{x}^{\,l}. \tag{81}$$

This completes the proof. □

Going back to Eq. 22 of Main Text, we find as a consequence of Claim 5.1 that

$$\bigoplus_{k=1}^{K}\bigoplus_{q=1}^{Q_k}\vec{x}_{kq} = \bigoplus_{k=1}^{K}\bigoplus_{q=1}^{Q_k} c_{qk}\vec{x}_k = \left[\bigoplus_{k=1}^{K}\bigoplus_{q=1}^{Q_k} c_{qk} I_{M_k\times M_k}\right]\left[\bigoplus_{k=1}^{K}\bigoplus_{q=1}^{Q_k}\vec{x}_k\right] \simeq \bigoplus_{k=1}^{K}\bigoplus_{q=1}^{Q_k}\vec{x}_k = \bigoplus_{k=1}^{K} Q_k\vec{x}_k, \tag{82}$$

where $I_{M_k\times M_k}$ denotes the identity matrix. Note that the penultimate step follows from the fact that multiplication by the matrix $\bigoplus_{k=1}^{K}\bigoplus_{q=1}^{Q_k} c_{qk} I_{M_k\times M_k}$ is a similarity transformation. Eq. 22 of Main Text results.

#### 5.2 Generalized distillations of $H_{\text{PSCP}}$-equivariant embeddings

Here we remove the assumption in the Main Text that every irreducible embedding $\vec{x}_{kq}$ in Eq. 16 is nonzero. Removing this assumption and retracing the logic, it is straight-forward to see that only two changes to the Main Text equations are required. First, Eq. 12 of Main Text becomes

$$\vec{x}^{\,\text{dist}} = \bigoplus_{k\in\mathcal{K}}\vec{x}_k, \tag{83}$$

where $\mathcal{K}\subseteq\mathcal{I}_K$ is the set of $k$ such that $\vec{x}_{qk}\neq\vec{0}$ for at least one value of $q$, and where $\vec{x}_k$ is chosen to be any one of the nonzero embeddings $\vec{x}_{qk}$. Second, the statement that the number of gauge freedoms equals the sum of the degrees of all redundant irreducible representations still holds, but the term "redundant" requires clarification. For every $k\in\mathcal{I}_K$, there are $Q_k$ copies of the representation $R_k$ in the Maschke decomposition (Eq. 15 of Main Text). When $k\in\mathcal{K}$, $Q_k-1$ of these copies of $R_k$ are redundant, because all but one of the $Q_k$ embeddings $\vec{x}_{kq}$ can be zeroed out by similarity transformations. When $k\in\mathcal{I}_K\setminus\mathcal{K}$, however, all $Q_k$ of these copies of $R_k$ are redundant because all of the $\vec{x}_{kq}$ are already zero. Eq. 22 of Main Text therefore becomes

$$\gamma \;=\; \dim\vec{x}-\dim\vec{x}^{\,\text{dist}} \;=\; \sum_{k\in\mathcal{K}}(Q_k-1)\deg R_k \;+\sum_{k\in\mathcal{I}_K\setminus\mathcal{K}} Q_k\deg R_k. \tag{84}$$

#### 5.3 Restriction to the reals

**Claim 5.2.** *If an embedding $\vec{x}$ is real then there is a corresponding real distillation matrix $T_{\text{dist}}$.*

*Proof.* Given an embedding $\vec{x}$, the embedding distillation procedure yields a distillation matrix $T_{\text{dist}}$ such that

$$T_{\text{dist}}\,\vec{x} = \vec{x}^{\,\text{dist}}\oplus\vec{0}_\gamma. \tag{85}$$

If $\vec{x}$ is restricted to the reals, and we construct $\vec{x}^{\,\text{dist}}$ using the choices of $\vec{x}^{\,\text{triv}}$ and $\vec{x}^{\,\text{sim}}$ described in this work (both of which are real), then taking one half the sum of Eq. 86 and its complex conjugate yields

$$\Re(T_{\text{dist}})\vec{x} = \vec{x}^{\,\text{dist}}\oplus\vec{0}_\gamma. \tag{86}$$

where $\Re(T_{\text{dist}})$ denotes the real part of $T_{\text{dist}}$. We therefore find that $\Re(T_{\text{dist}})$ is an equivalent real distillation matrix. In particular, the last $\gamma$ rows of $T_{\text{dist}}$ provide a real basis for the space of gauge freedoms $G$. We conclude that our embedding distillation procedure works when $\vec{x}$ and $\vec{\theta}$ are restricted to the reals. □

### 6 Analytic results for specific generalized one-hot models (Table 1 of Main Text)

We now compute the number of parameters and gauge freedoms for a variety of generalized one-hot models. To avoid conflicts in notation, we use $N$ and $n$ in place of $K$ and $k$ to parameterize orders of interactions. Note that $N$ here does not refer the number of sequences in $\mathcal{S}$.

#### 6.1 $N$-order model

Here we compute the number of parameters and gauge freedoms for the $N$-order model. We begin by computing the dimension of the model embedding $\vec{x}_{N\text{-order}}$. The position sets that define $\vec{x}_{N\text{-order}}$ are

$$\{A_j\}_{j=1}^{J} = \{U : U\subseteq I_L, |U| = N\}. \tag{87}$$

From this we see that there are $J = \binom{L}{N}$ distinct sets $A_j$, and that $|A_j| = N$ for every $j \in \mathcal{I}_J$. By Main Text Eq. 30, the dimension of $\vec{x}_{N\text{-order}}$ is therefore

$$\dim \vec{x}_{N\text{-order}} = \sum_{j=1}^{J} \alpha^{|A_j|} \tag{88}$$

$$= \binom{L}{N} \alpha^N. \tag{89}$$

Next we compute the dimension of the distilled embedding $\dim \vec{x}^{\text{dist}}_{N\text{-order}}$. The position sets that define $\vec{x}^{\text{dist}}_{N\text{-order}}$ are

$$\{B_k\}_{k=1}^{K} = \{V : V \subseteq A_j, j \in \mathcal{I}_J\} \tag{90}$$

$$= \{V : V \subseteq U, U \subseteq I_L, |U| = N\} \tag{91}$$

$$= \{V : V \subseteq I_L, |V| \leq N\} \tag{92}$$

$$= \bigcup_{n=0}^{N} \{V : V \subseteq I_L, |V| = n\}, \tag{93}$$

where the union is over disjoint sets. From this we see that, for each $n \in \{0, \ldots, N\}$, there are $\binom{L}{n}$ sets $B_k$ such that $|B_k| = n$. By Main Text Eq. 32, the dimension of $\vec{x}^{\text{dist}}_{N\text{-order}}$ is therefore

$$\dim \vec{x}^{\text{dist}}_{N\text{-order}} = \sum_{k=1}^{K} (\alpha - 1)^{|B_k|} \tag{94}$$

$$= \sum_{n=0}^{N} \binom{L}{n} (\alpha - 1)^n. \tag{95}$$

The number of parameters, $M_{N\text{-order}}$, and the number of gauge freedoms, $\gamma_{N\text{-order}}$, are therefore given by

$$M_{N\text{-order}} = \dim \vec{x}_{N\text{-order}} \tag{96}$$

$$= \binom{L}{N} \alpha^N, \tag{97}$$

$$\gamma_{N\text{-order}} = \dim \vec{x}_{N\text{-order}} - \dim \vec{x}^{\text{dist}}_{N\text{-order}} \tag{98}$$

$$= \binom{L}{N} \alpha^N - \sum_{n=0}^{N} \binom{L}{n} (\alpha - 1)^n. \tag{99}$$

#### 6.2 Hierarchical $N$-order models

We define the hierarchical $N$-order model to be the sum of $n$-order models taken over all $n \in \{0, 1, \ldots, N\}$. The embedding of the hierarchical $N$-order model is

$$\vec{x}_{\leq N\text{-order}} = \bigoplus_{n=0}^{N} \vec{x}_{n\text{-order}}. \tag{100}$$

The number of parameters of the hierarchical $N$-order model is therefore

$$M_{\leq N\text{-order}} = \sum_{n=0}^{N} M_{n\text{-order}} \tag{101}$$

$$= \sum_{n=0}^{N} \binom{L}{n} \alpha^n. \tag{102}$$

Since every parameter of the models of order $N-1$ and lower is redundant with parameters of the model of order $N$, the number of gauge freedoms of the hierarchical $N$-order model is

$$\gamma_{\leq N\text{-order}} = \dim \vec{x}_{\leq N\text{-order}} - \dim \vec{x}^{\text{dist}}_{\leq N\text{-order}} \tag{103}$$

$$= M_{\leq N\text{-order}} - \dim \vec{x}^{\text{dist}}_{N\text{-order}} \tag{104}$$

$$= \sum_{n=0}^{N} \binom{L}{n} \alpha^n - \sum_{n=0}^{N} \binom{L}{n} (\alpha - 1)^n \tag{105}$$

$$= \sum_{n=0}^{N} \binom{L}{n} [\alpha^n - (\alpha - 1)^n]. \tag{106}$$

Additive models, pairwise models, and all-order models are special cases of the hierarchical $N$-order model.

###### 6.2.1 $N = 1$: Additive model

The additive model is a hierarchical 1-order model. Setting $N = 1$, Eq. 102 gives

$$M_{\text{additive}} = M_{\leq 1\text{-order}} \tag{107}$$
$$= \binom{L}{0} + \binom{L}{1}\alpha \tag{108}$$
$$= 1 + L\alpha, \tag{109}$$

and Eq. 106 gives,

$$\gamma_{\text{additive}} = \gamma_{\leq 1\text{-order}}, \tag{110}$$
$$= \binom{L}{0}(1-1) + \binom{L}{1}(\alpha - (\alpha - 1)), \tag{111}$$
$$= L. \tag{112}$$

###### 6.2.2 $N = 2$: Pairwise model

The pairwise model is a hierarchical 2-order model. Setting $N = 2$, Eq. 102 gives

$$M_{\text{pairwise}} = M_{\leq 2\text{-order}} \tag{113}$$
$$= \binom{L}{0} + \binom{L}{1}\alpha + \binom{L}{2}\alpha^2 \tag{114}$$
$$= 1 + L\alpha + \binom{L}{2}\alpha^2, \tag{115}$$

and Eq. 106 gives

$$\gamma_{\text{pairwise}} = \gamma_{\leq 2\text{-order}} \tag{116}$$
$$= \binom{L}{0}(1-1) + \binom{L}{1}(\alpha - (\alpha-1)) + \binom{L}{2}(\alpha^2 - (\alpha-1)^2) \tag{117}$$
$$= L + \binom{L}{2}(2\alpha - 1). \tag{118}$$

###### 6.2.3 $N = L$: All-order model

The all-order model is a hierarchical $L$-order model. Setting $N = L$, Eq. 102 gives

$$M_{\text{all-order}} = M_{\leq L\text{-order}} \tag{119}$$
$$= \sum_{n=0}^{L} \binom{L}{n}\alpha^n \tag{120}$$
$$= (\alpha + 1)^L, \tag{121}$$

and Eq. 106 gives,

$$\gamma_{\text{all-order}} = \gamma_{\leq L\text{-order}}, \tag{122}$$
$$= \sum_{n=0}^{L} \binom{L}{n} \left[\alpha^n - (\alpha - 1)^n\right], \tag{123}$$
$$= (\alpha + 1)^L - \alpha^L. \tag{124}$$

##### 6.3 $N$-adjacent model

Here we compute the number of parameters and gauge freedoms for the $N$-adjacent model. To facilitate computations, we define $\operatorname{span} U = \max(U) - \min(U) + 1$ for any nonempty set $U \subseteq \mathcal{I}_l$, as well as $\operatorname{span} \emptyset = 0$. We also define the 0-adjacent model to be the constant model, i.e., the model that corresponds to the embedding $\vec{x}_{\text{constant}} = \vec{x}^{\,\text{triv}}$, which has $M_{\text{constant}} = 1$ parameters and $\gamma_{\text{constant}} = 0$ gauge freedoms.

For $N \geq 1$, we begin by computing the degree of the embedding $\vec{x}_{N\text{-adjacent}}$. The position sets that define $\vec{x}_{N\text{-adjacent}}$ are

$$\begin{aligned} \{A_j\}_{j=1}^{J} &= \{U : U \subseteq I_L,\ |U| = N,\ \text{span}\,(U) = N\} && (125)\\ &= \{U_l : l \in \mathcal{I}_{L-N+1}\}, \quad \text{where} \quad U_l = \{l, \ldots, l+N-1\}. && (126)\end{aligned}$$

From this we see that there are $J = L - N + 1$ distinct sets $A_j$, and that $|A_j| = N$ for every $j \in \mathcal{I}_J$. By Main Text Eq. 30, the dimension of $\vec{x}_{N\text{-adjacent}}$ is therefore

$$\begin{aligned} \dim \vec{x}_{N\text{-adjacent}} &= \sum_{j=1}^{J} \alpha^{|A_j|} && (127)\\ &= (L-N+1)\alpha^{N}. && (128)\end{aligned}$$

Next we compute the dimension of the distilled embedding $\vec{x}^{\,\text{dist}}_{N\text{-adjacent}}$. The position sets that define $\vec{x}^{\,\text{dist}}_{N\text{-adjacent}}$ are

$$\begin{aligned} \{B_k\}_{k=1}^{K} &= \{V : V \subseteq A_j,\ j \in \mathcal{I}_J\} && (129)\\ &= \{V : V \subseteq U,\ U \subseteq I_L,\ |U| = N,\ \text{span}\,(U) = N\} && (130)\\ &= \{V : V \subseteq I_L,\ |V| \leq N,\ \text{span}\,(V) \leq N\} && (131)\\ &= \bigcup_{p=0}^{N} \bigcup_{n=0}^{p} \{V : V \subseteq I_L,\ |V| = n,\ \text{span}\,(V) = p\}, && (132)\end{aligned}$$

where the unions are over disjoint sets, and we have used the fact that $|V| \leq \text{span}\,(V)$ for all $V \subseteq \mathcal{I}_L$. By Main Text Eq. 32, the dimension of $\vec{x}^{\,\text{dist}}$ is therefore

$$\begin{aligned} \dim \vec{x}^{\,\text{dist}}_{N\text{-adjacent}} &= \sum_{k=1}^{K} (\alpha-1)^{|B_k|} && (133)\\ &= \sum_{p=0}^{N} \sum_{n=0}^{p} z(n,p)(\alpha-1)^{n}, && (134)\end{aligned}$$

where $z(n,p)$ is the number of subsets of $\mathcal{I}_L$ that have $n$ elements and span $p$.

- For $n = 0$, only $V = \emptyset$ has span $p = 0$. Therefore $z(0,0) = 1$, and $z(0,p) = 0$ for all $p \neq 0$.
- For $n = 1$, there are $L$ subsets: $U = \{l\}$ for every $l \in \mathcal{I}_L$. Each of these subsets has span $p = 1$. Therefore $z(1,1) = L$ and $z(1,p) = 0$ for all $p \neq 1$.
- For every $n \in \{2, \ldots, N\}$, there is at least one subset $U$ having span $p$ for every $p \in \{2, \ldots, N\}$. Given the endpoints of $U$, there are $\binom{p-2}{n-2}$ ways to choose the elements of $U$. And given $p$, there are $L - p + 1$ possible choices for the endpoints of $U$. Therefore, $z(n,p) = (L-p+1)\binom{p-2}{n-2}$.

Continuing from Eq. 134,

$$\begin{aligned} \dim \vec{x}^{\,\text{dist}}_{N\text{-adjacent}} &= 1 + L(\alpha-1) + \sum_{p=2}^{N} \sum_{n=2}^{N} \binom{p-2}{n-2} (L-p+1)(\alpha-1)^{n} && (135)\\ &= 1 + L(\alpha-1) + (\alpha-1)^2 \sum_{p=2}^{N} (L-p+1)\alpha^{p-2} && (136)\\ &= 1 + L(\alpha-1) + (\alpha-1)^2 (L-1) \sum_{m=0}^{N-2} \alpha^{m} - (\alpha-1)^2 \sum_{m=0}^{N-2} m\alpha^{m}. && (137)\end{aligned}$$

Setting $M = N - 2$, and using

$$\sum_{m=0}^{M} \alpha^{m} = \frac{\alpha^{M+1} - 1}{\alpha - 1}, \qquad (138)$$

and

$$\sum_{m=0}^{M} m\alpha^{m} = \alpha \frac{d}{d\alpha} \sum_{m=0}^{M} \alpha^{m} = \frac{\alpha}{(\alpha-1)^2} \left[1 - \alpha^{M+1} + (M+1)(\alpha-1)\alpha^{M}\right], \qquad (139)$$

we get (after a remarkable simplification),

$$\dim \vec{x}^{\text{dist}}_{N\text{-adjacent}} = (L-N+1)\alpha^N - (L-N)\alpha^{N-1}. \tag{140}$$

The number of parameters, $M_{N\text{-adjacent}}$, and the number of gauge freedoms, $\gamma_{N\text{-adjacent}}$, is (for $N \geq 1$) therefore given by

$$\begin{aligned} M_{N\text{-adjacent}} &= \dim \vec{x}_{N\text{-adjacent}}, & (141)\\ &= (L-N+1)\alpha^N, & (142)\\ \gamma_{N\text{-adjacent}} &= \dim \vec{x}_{N\text{-adjacent}} - \dim \vec{x}^{\text{dist}}_{N\text{-adjacent}}, & (143)\\ &= (L-N)\alpha^{N-1}. & (144) \end{aligned}$$

Note the curious fact that the number of gauge freedoms for the $N$-adjacent model is equal to the number of parameters for the $(N-1)$-adjacent model, i.e.

$$\gamma_{N\text{-adjacent}} = M_{(N-1)\text{-adjacent}}. \tag{145}$$

#### 6.4 Hierarchical $N$-adjacent models

We define the hierarchical $N$-adjacent model to be the sum of $n$-adjacent terms taken over all $n \in \{0, 1, \dots, N\}$. The embedding of the hierarchical $N$-adjacent model is

$$\vec{x}_{\leq N\text{-adjacent}} = \bigoplus_{n=0}^{N} \vec{x}_{n\text{-adjacent}}. \tag{146}$$

The number of parameters of the hierarchical $N$-adjacent model is therefore,

$$\begin{aligned} M_{\leq N\text{-adjacent}} &= \sum_{n=0}^{N} M_{n\text{-adjacent}}, & (147)\\ &= 1 + \sum_{n=1}^{N} (L-n+1)\alpha^n, & (148)\\ &= 1 + \frac{\alpha}{(\alpha-1)^2}\left[(L-N+1)\alpha^{N+1} - (L-N)\alpha^N - (L+1)\alpha + L\right]. & (149) \end{aligned}$$

Since every parameter of the models of order $N-1$ and lower is redundant with parameters of the model of order $N$, the number of gauge freedoms of the hierarchical $N$-adjacent model,

$$\begin{aligned} \gamma_{\leq\, N\text{-adjacant}} &= \dim \vec{x}_{\leq N\text{-adjacent}} - \dim \vec{x}^{\text{dist}}_{\leq N\text{-adjacent}} & (150)\\ &= M_{\leq N\text{-adjacent}} - \dim \vec{x}^{\text{dist}}_{N\text{-adjacent}} & (151)\\ &= 1 + \frac{\alpha}{(\alpha-1)^2}\left[(L-N+1)\alpha^{N+1} - (L-N)\alpha^N - (L+1)\alpha + L\right] & (152)\\ &\quad -(L-N+1)\alpha^N + (L-N)\alpha^{N-1}, & (153)\\ &= (L-N)\alpha^{N-1} + 1 + \frac{\alpha}{(\alpha-1)^2}\left[(L-N+2)\alpha^N - (L-N+1)\alpha^{N-1} - (L+1)\alpha + L\right]. & (154) \end{aligned}$$

Additive models, nearest-neighbor models, and all-order models are special cases of the hierarchical $N$-adjacent model.

##### 6.4.1 $N = 1$: Additive model

The additive model is a hierarchical 1-adjacent odel. Setting $N = 1$, Eq. 149 gives

$$\begin{aligned} M_{\text{additive}} &= M_{\leq 1\text{-adjacent}} & (155)\\ &= 1 + \frac{\alpha}{(\alpha-1)^2}\left[L\alpha^2 - (L-1)\alpha - (L+1)\alpha + L\right] & (156)\\ &= 1 + L\alpha, & (157) \end{aligned}$$

and Eq. 154 gives

$$\begin{aligned} \gamma_{\text{additive}} &= \gamma_{\leq 1\text{-adjacent}} & (158)\\ &= (L-1) + 1 + \frac{\alpha}{(\alpha-1)^2}\left[(L+1)\alpha - L - (L+1)\alpha + L\right] & (159)\\ &= L. & (160) \end{aligned}$$

##### 6.4.2 $N = 2$: Nearest-neighbor model

The nearest-neighbor model is a hierarchical 2-adjacent model. Setting $N = 2$, Eq. 149 gives

$$\begin{aligned} M_{\text{neighbor}} &= M_{\leq 2\text{-adjacent}} && (161)\\ &= 1 + \frac{\alpha}{(\alpha-1)^2}\left[(L-1)\alpha^3 - (L-2)\alpha^2 - (L+1)\alpha + L\right] && (162)\\ &= 1 + L\alpha + (L-1)\alpha^2, && (163)\end{aligned}$$

and Eq. 154 gives

$$\begin{aligned} \gamma_{\text{neighbor}} &= \gamma_{\leq 2\text{-adjacent}} && (164)\\ &= (L-2)\alpha + 1 + \frac{\alpha}{(\alpha-1)}\left[L\alpha^2 - (L-1)\alpha - (L+1)\alpha + L\right] && (165)\\ &= 1 + 2(L-1)\alpha. && (166)\end{aligned}$$

##### 6.4.3 $N = L$: All-adjacent model

The all-order adjacent model is a hierarchical $L$-adjacent model. Setting $N = L$, Eq. 149 gives

$$\begin{aligned} M_{\text{all-adjacent}} &= M_{\leq L\text{-adjacent}} && (167)\\ &= 1 + \frac{\alpha}{(\alpha-1)^2}\left[\alpha^{L+1} - (L+1)\alpha + L\right], && (168)\end{aligned}$$

and Eq. 154 gives,

$$\begin{aligned} \gamma_{\text{all-adjacent}} &= \gamma_{\leq L\text{-adjacent}} && (169)\\ &= 1 + \frac{\alpha}{(\alpha-1)^2}\left[2\alpha^{L} - \alpha^{L-1} - (L+1)\alpha + L\right]. && (170)\end{aligned}$$

### 7 Results for other symmetry groups: $H_{\text{GCP}}$, $H_{\text{PP}}$, and $H_{\text{PSCP}}$

#### 7.1 Embedding distillation does not work for $H_{\text{GCP}}$ or $H_{\text{PP}}$

Embedding distillation does not work for $H_{\text{GCP}}$ (the group of global character permutations) or for $H_{\text{PP}}$ (the group of position permutations). The reason is that Claim 5.1 (Theorem 1 of Main Text), does not hold for either of these groups: two embeddings that transform under the same representation are not necessarily proportional (or, in fact, related by any similarity transformation). Consider, for example, the following one-dimensional embeddings of a length $L$ sequence:

$$\vec{x}_1(s) = \begin{cases} [1] \text{ if } s_l = s'_l \text{ for all } l, l' \in \mathcal{I}_L \\ [0] \text{ otherwise} \end{cases}, \quad (171)$$

$$\vec{x}_2(s) = \begin{cases} [0] \text{ if } s_l = s'_l \text{ for all } l, l' \in \mathcal{I}_L \\ [1] \text{ otherwise} \end{cases}. \quad (172)$$

Both $\vec{x}_1$ and $\vec{x}_2$ are invariant under $H_{\text{GCP}}$ and $H_{\text{PP}}$, and therefore transform under the trivial representation of both groups. However, $\vec{x}_1$ and $\vec{x}_2$ are not related by any similarity transformation, as no similarity transformation can map a zero vector to a nonzero vector, thus showing that Claim 5.1 does not hold for $H_{\text{GCP}} or H_{\text{PP}}$. Consequently, distillation cannot be used to determine the gauge freedoms of either embedding, and the number of gauge freedoms cannot be computed by summing the degrees of the redundant irreducible representations under which these embeddings transform.

To see explicitly how embedding distillation fails for $H_{\text{GCP}}$ and for $H_{\text{PP}}$, consider the additive embedding:

$$\vec{x}_{\text{additive}} = \vec{x}^{\,\text{triv}} \oplus \bigoplus_{l=1}^{L} \vec{x}_l^{\,\text{ohe}}. \quad (173)$$

Under $H_{\text{GCP}}$, each $\vec{x}_l^{\,\text{ohe}}$ transforms under the same representation: $R^{\text{ohe}}_{(\alpha)}$, the one-hot representation of $S_\alpha$ (we indicate the degree of the representation in the subscript for clarity). Using $R^{\text{ohe}}_{(\alpha)} \simeq R^{\text{triv}} \oplus R^{\text{sim}}_{(\alpha-1)}$, where $R^{\text{sim}}_{(\alpha-1)}$ is the simplex representation of $S_\alpha$, we find that $\vec{x}_{\text{additive}}$ transforms under $H_{\text{GCP}}$ according to the representation

$$\begin{aligned} R^{\text{additive}}_{\text{GCP}} &= R^{\text{triv}} \oplus [I_L \otimes R^{\text{ohe}}_{(\alpha)}], && (174)\\ &\simeq R^{\text{triv}} \oplus L R^{\text{ohe}}_{(\alpha)}, && (175)\\ &\simeq R^{\text{triv}} \oplus L[R^{\text{triv}} \oplus R^{\text{sim}}_{(\alpha-1)}], && (176)\\ &\simeq [L+1]R^{\text{triv}} \oplus L R^{\text{sim}}_{(\alpha-1)}, && (177)\\ &\simeq R^{\text{dist}}_{\text{GCP}} \oplus R^{\text{redun}}_{\text{GCP}}, && (178)\end{aligned}$$

where the distilled representation is

$$R_{\mathrm{GCP}}^{\mathrm{dist}} = R^{\mathrm{triv}} \oplus R_{(\alpha-1)}^{\mathrm{sim}}, \tag{179}$$

and the redundant irreducible representations are collected into

$$R_{\mathrm{GCP}}^{\mathrm{redun}} = LR^{\mathrm{triv}} \oplus (L-1)R_{(\alpha-1)}^{\mathrm{sim}}. \tag{180}$$

We thus see that

$$\deg R_{\mathrm{GCP}}^{\mathrm{redun}} = \alpha L - \alpha + 1. \tag{181}$$

This does not match the number of gauge freedoms, $\gamma_{\mathrm{additive}} = L$. The reason is that each copy of $R_{(\alpha-1)}^{\mathrm{sim}}$ transforms a simplex embedding of $\mathcal{A}$, one for each position in $\mathcal{I}_L$, and these embeddings are not equivalent. Similarly, $H_{\mathrm{PP}}$ acts on $\vec{x}_{\mathrm{additive}}$ via the representation

$$\begin{aligned} R_{\mathrm{PP}}^{\mathrm{additive}} &= R^{\mathrm{triv}} \oplus [R_{(L)}^{\mathrm{ohe}} \otimes I_\alpha], & (182)\\ &= R^{\mathrm{triv}} \oplus \alpha R_{(L)}^{\mathrm{ohe}}, & (183)\\ &\simeq R^{\mathrm{triv}} \oplus \alpha[R^{\mathrm{triv}} \oplus R_{(L-1)}^{\mathrm{sim}}], & (184)\\ &\simeq [1+\alpha]R^{\mathrm{triv}} \oplus \alpha R_{(L-1)}^{\mathrm{sim}}, & (185)\\ &\simeq R_{\mathrm{PP}}^{\mathrm{dist}} \oplus R_{\mathrm{PP}}^{\mathrm{redun}}, & (186) \end{aligned}$$

where the distilled representation is

$$R^{\mathrm{dist}} = R^{\mathrm{triv}} \oplus R_{(L-1)}^{\mathrm{sim}}, \tag{187}$$

and the redundant irreducible representations are collected into

$$R_{\mathrm{PP}}^{\mathrm{redun}} = \alpha R^{\mathrm{triv}} \oplus (\alpha-1)R_{(L-1)}^{\mathrm{sim}}. \tag{188}$$

We thus see that

$$\deg R_{\mathrm{PP}}^{\mathrm{redun}} = \alpha L - L + 1. \tag{189}$$

This does not match the number of gauge freedoms $\gamma_{\mathrm{additive}} = L$. The reason is that each copy of $R_{(L-1)}^{\mathrm{sim}}$ transforms a simplex embedding of $\mathcal{I}_L$, one for each character in $\mathcal{A}$, and these embeddings are not equivalent.

#### 7.2 Embedding distillation does work for $H_{\mathrm{Ham}}$

Our embedding distillation procedure does work for the symmetry group of the Hamming graph. This group, which we denote by $H_{\mathrm{Ham}}$, is given by the semidirect product

$$H_{\mathrm{Ham}} = H_{\mathrm{PSCP}} \rtimes H_{\mathrm{PP}}. \tag{190}$$

From this definition we see that $H_{\mathrm{PSCP}}$ is a normal subgroup of $H_{\mathrm{Ham}}$.

**Claim 7.1.** *Nonzero sequence embeddings that transform under the same irreducible representation of $H_{\mathrm{Ham}}$ are equivalent.*

*Proof.* Assume $\vec{x}$ and $\vec{y}$ are two nonzero embeddings that transform under the same irreducible representation $R$ of $H_{\mathrm{Ham}}$. Let $R'$ denote $R$ restricted to $H_{\mathrm{PSCP}}$. By Maschke's theorem, there is a similarity transformation matrix $T_{\mathrm{dist}}$ such that

$$R' = T\left[\bigoplus_{k=1}^{K} R'_k\right]T^{-1}, \tag{191}$$

where, for each $k = 1, \ldots, K$, $R'_k$ is an irreducible representation of $H_{\mathrm{PSCP}}$. Consequently,

$$\vec{x} = T\left[\bigoplus_{k=1}^{K} \vec{x}_k\right], \quad \text{and} \quad \vec{y} = T\left[\bigoplus_{k=1}^{K} \vec{y}_k\right], \tag{192}$$

where, for each $k \in \mathcal{I}_K$, both $\vec{x}_k$ and $\vec{y}_k$ transform under $R'_k$. We now show that all $\vec{x}_k$ and $\vec{y}_k$ are nonzero. Let $m_k = \deg R'_k$, $\mathcal{K} = \left\{k : k \in \mathcal{I}_K,\ \vec{x}_k = \vec{0}\right\}$, and define the vector space

$$V = T^{-1\dagger}\left[\bigoplus_{k=1}^{K} \begin{cases} \mathbb{R}^{m_k} & \text{if } k \in \mathcal{K}, \\ \left\{\vec{0}\right\} & \text{otherwise} \end{cases}\right]. \tag{193}$$

It is readily see that $V$ is orthogonal to span $\vec{x}$, i.e., $\vec{v}^\dagger \vec{x}(s) = 0$ for all $v \in V$ and all $s \in \mathcal{S}$. But we know from Claim 3.3 that, because $\vec{x}$ transforms under an irreducible representation of $H_{\text{Ham}}$, span $\vec{x}$ has full rank. Consequently, $\mathcal{K} = \emptyset$, and thus all $\vec{x}_k$ are nonzero. Similarly, all $\vec{y}_k$ are nonzero. Since both $\vec{x}_k$ and $\vec{y}_k$ are nonzero embeddings that transform under the same irreducible representation of $H_{\text{PSCP}}$, Claim 5.1 tells us that $\vec{y}_k = c_k \vec{x}_k$ for some nonzero scalar $c_k \in \mathbb{R}$. Therefore,

$$\vec{y} = TCT^{-1}\vec{x} \quad \text{where} \quad C = \bigoplus_{k=1}^{K} c_k I_{m_k}. \tag{194}$$

This shows that $\vec{x}$ and $\vec{y}$ are related by a similarity transformation, thereby completing the proof. □

### 8 Distillation algorithm

We now describe the embedding distillation algorithm discussed in the section "Computational analysis of models" of Main Text and illustrated in main text Fig. 4. Python code implementing this algorithm is available at https://github.com/jbkinney/23_posfai.

#### 8.1 Overview of the algorithm

Let $\vec{x}$ be an equivariant one-hot embedding of sequences as defined in Main Text. Specifically, define

$$\vec{x} = \bigoplus_{j=1}^{J} \bigotimes_{l \in A_j} \vec{x}_l^{\,\text{ohe}}, \tag{195}$$

where each $A_j$, $j \in \mathcal{I}_J$, denotes an ordered set of sequence positions, and $A = (A_j)_{j=1}^{J}$ denotes an ordered set of position subsets the sets of positions used to construct the embedding. Distillation consists of multiplying $\vec{x}$ by a "distillation matrix", $T_{\text{dist}}$, such that

$$T_{\text{dist}}\,\vec{x} = \vec{x}^{\,\text{dist}} \oplus \vec{0}_\gamma, \tag{196}$$

where $\vec{x}^{\,\text{dist}}$ is the full-rank distilled embedding, $\vec{0}_\gamma$ is a vector of $\gamma$ zeros, and $\gamma$ is the number of gauge freedoms of $\vec{x}$. As in Main Text, we construct $T_{\text{dist}}$ by expressing it as

$$T_{\text{dist}} = T_{\text{sort}}\,T_{\text{thin}}\,T_{\text{decom}}. \tag{197}$$

We now describe the operations each of the three factors above perform:

1. $T_{\text{decom}}$ decomposes each Kronecker product of $\vec{x}_l^{\,\text{ohe}}$ into a direct sum of Kronecker products of $\vec{x}_l^{\,\text{sim}}$. Specifically, multiplying Eq. 195 by $T_{\text{decom}}$ yields,

$$T_{\text{decom}}\,\vec{x} \quad = \quad \bigoplus_{p=1}^{P} \vec{x}_{k_p}, \tag{198}$$

   where each $\vec{x}_k$ is an irreducible representation given by

$$\vec{x}_k = \bigotimes_{l \in B_k} \vec{x}_l^{\,\text{sim}}, \tag{199}$$

   each $B_k$ is a distinct ordered set of positions (in which positions are listed in increasing order), $B = (B_k)_{k=1}^{K}$ is an ordered set comprising all unique subsets of positions found within all the $A_j$, and $(k_p)_{p=1}^{P}$ is an ordered set of (generally non-unique) indices within $\mathcal{I}_K$.

2. $T_{\text{thin}}$ thins the resulting embedding by zeroing-out all redundant copies of each irreducible embedding. Specifically, multiplying Eq. 198 by $T_{\text{thin}}$ yields

$$T_{\text{thin}}\,T_{\text{decom}}\,\vec{x} = \bigoplus_{p=1}^{P} \begin{cases} \vec{x}_{k_p} & \text{if } k_p \neq k_{p'} \text{ for any } p' < p \\ \vec{0}_{m_{k_p}} & \text{otherwise} \end{cases}, \tag{200}$$

   where each $m_k = (\alpha - 1)^{|B_k|}$ is the degree of $\vec{x}_k$.

3. $T_{\text{sort}}$ sorts the embeddings in the direct sum so that all nonzero embeddings come first. Specifically, multiplying Eq. 200 by $T_{\text{sort}}$ yields

$$T_{\text{sort}}\, T_{\text{thin}}\, T_{\text{decom}}\, \vec{x} = \vec{x}^{\,\text{dist}} \oplus \vec{0}_{\gamma}, \tag{201}$$

where

$$\vec{x}^{\,\text{dist}} = \bigoplus_{k=1}^{K} \vec{x}_k. \tag{202}$$

We now show how to compute each of these three matrix factors. We find that the resulting distillation matrix $T_{\text{dist}}$ is sparse (because $T_{\text{decom}}$, $T_{\text{thin}}$, and $T_{\text{sort}}$ are sparse) and that all nonzero entries of $T_{\text{dist}}$ are either $+1$ or $-1$. We also show how to compute the inverse of each factor without having to invert any matrices, thereby allowing the rapid computation of

$$T_{\text{dist}}^{-1} = T_{\text{decom}}^{-1}\, T_{\text{thin}}^{-1}\, T_{\text{sort}}^{-1}. \tag{203}$$

Note: In what follows we index vector and matrix elements starting from zero. This differs from the one-indexing used in Main Text and elsewhere in the supplement. We adopt this indexing change here to aid coding efforts.

#### 8.2 Computation of the decomposition matrix $T_{\text{decom}}$.

We build the decomposition matrix, $T_{\text{decom}}$, as a direct sum of individual $N$th order decomposition matrices, $T_{\text{decom}}^{(N)}$. Specifically,

$$T_{\text{decom}} = \bigoplus_{j=1}^{J} T_{\text{decom}}^{(N_j)}, \tag{204}$$

where $N_j = |A_j|$ for all $j \in \mathcal{I}_J$. Given $N$ we define $T_{\text{decom}}^{(N)}$ so that

$$T_{\text{decom}}^{(N)} \left[ \bigotimes_{n=1}^{N} \vec{x}_{l_n}^{\,\text{ohe}} \right] = \bigoplus_{B \subseteq A} \left[ \bigotimes_{l \in B} \vec{x}_{l}^{\,\text{sim}} \right] \tag{205}$$

for any subset of positions $A$ that has $N$ elements. We computationally construct $T_{\text{decom}}^{(N)}$ by recursion using

$$T_{\text{decom}}^{(N)} = T_{\text{perm}}^{(N)} \left[ T_{\text{decom}}^{(N-1)} \otimes T \right] \tag{206}$$

where $T$ is the single-position decomposition matrix. Using this recursion relation requires defining $T$ and $T_{\text{perm}}^{(N)}$:

- $T$ is an $\alpha \times \alpha$ matrix having elements

$$[T]_{ij} = \begin{cases} 1 & \text{if } i = 0, \\ -1 & \text{if } 0 < i < \alpha,\ j = \alpha - 1, \\ 1 & \text{if } 0 < i < \alpha,\ j = i - 1, \\ 0 & \text{otherwise}, \end{cases} \tag{207}$$

  for $i, j \in \{0, \ldots, \alpha - 1\}$. Note that $T$ matches Eq. 20 when $\alpha = 4$.

- $T_{\text{perm}}^{(N)}$ is an $\alpha^N \times \alpha^N$ permutation matrix having elements

$$\left[ T_{\text{perm}}^{(N)} \right]_{ij} = \begin{cases} 1 & \text{if } 0 \leq i < \alpha^{N-1} \text{ and } \ j = \alpha i, \\ 1 & \text{if } \alpha^{N-1} \leq i < \alpha^{N} \text{ and } j = 1 + (i - \alpha^{N-1}) + \lfloor (i - \alpha^{N-1})/(\alpha - 1) \rfloor, \\ 0 & \text{otherwise}. \end{cases} \tag{208}$$

  for $i, j \in \{0, \ldots, \alpha^N - 1\}$. $T_{\text{perm}}$ needed because Kronecker products do not distribute over direct sums. Specifically, for a general vector $\vec{v}$ having $\alpha^N$ elements,

$$\vec{v} \otimes \left( \vec{x}^{\,\text{triv}} \oplus \vec{x}_l^{\,\text{sim}} \right) \neq \left( \vec{v} \otimes \vec{x}^{\,\text{triv}} \right) \oplus \left( \vec{v} \otimes \vec{x}_l^{\,\text{sim}} \right). \tag{209}$$

  The matrix $T_{\text{perm}}^{(N)}$ fixes this inequality, i.e.,

$$T_{\text{perm}} \left[ \vec{v} \otimes \left( \vec{x}^{\,\text{triv}} \oplus \vec{x}_l^{\,\text{sim}} \right) \right] = \left( \vec{v} \otimes \vec{x}^{\,\text{triv}} \right) \oplus \left( \vec{v} \otimes \vec{x}_l^{\,\text{sim}} \right). \tag{210}$$

Similarly, we build the inverse of the decomposition matrix, $T_{\text{decom}}^{-1}$, as a direct sum of $N$'th order inverse decomposition matrices $\left(T_{\text{decom}}^{(N)}\right)^{-1}$, which are computed by recursion using

$$\left(T_{\text{decom}}^{(N)}\right)^{-1} = \left[\left(T_{\text{decom}}^{(N-1)}\right)^{-1} \otimes T^{-1}\right] \left(T_{\text{perm}}^{(N)}\right)^{-1}, \tag{211}$$

with

$$\left[T^{-1}\right]_{ij} = \frac{1}{\alpha} \times \begin{cases} 1 & \text{if } j = 0, \\ (\alpha - 1) & \text{if } 0 < j < \alpha,\ i = j - 1, \\ -1 & \text{otherwise,} \end{cases} \tag{212}$$

for $i, j \in \{0, \ldots, \alpha - 1\}$. Note that $T^{-1}$ matches Eq. 20 when $\alpha = 4$. Also note that, because $T_{\text{perm}}^{(N)}$ is a permutation matrix,

$$\left(T_{\text{perm}}^{(N)}\right)^{-1} = \left(T_{\text{perm}}^{(N)}\right)^{\top}. \tag{213}$$

#### 8.3 Computation of the thinning matrix $T_{\text{thin}}$

For each $p \in \mathcal{I}_P$, define the index offset function

$$\text{offset}(p) = \sum_{p' < p} m_{k_{p'}}, \tag{214}$$

and the first occurence function

$$\text{first}(p) = \min\left\{p' : k_p' = k_p\right\}. \tag{215}$$

The thinning matrix, $T_{\text{thin}}$, is an $M \times M$ having elements

$$[T_{\text{thin}}]_{ij} = \begin{cases} 1 & \text{if } i = j, \\ -1 & \text{if } i \neq j,\ i = \text{offset}(p) + h, \text{ and } j = \text{offset}(\text{first}(p)) + h \text{ for some } p \in \mathcal{I}_P,\ h \in \left\{0, \ldots, m_{k_p} - 1\right\}, \\ 0 & \text{otherwise,} \end{cases} \tag{216}$$

for $i, j \in \{0, \ldots, M - 1\}$. One can readily verify that the inverse of the thinning matrix thus has elements

$$\left[T_{\text{thin}}^{-1}\right]_{ij} = \begin{cases} 1 & \text{if } i = j, \\ 1 & \text{if } i \neq j,\ i = \text{offset}(p) + h, \text{ and } j = \text{offset}(\text{first}(p)) + h \text{ for some } p \in \mathcal{I}_P,\ h \in \left\{0, \ldots, m_{k_p} - 1\right\}. \\ 0 & \text{otherwise.} \end{cases} \tag{217}$$

#### 8.4 Computation of the sorting matrix $T_{\text{sort}}$

We define the sorted index offset function as

$$\text{sortedoffset}(p) \quad = \quad \begin{cases} \text{offest}_1(p) & \text{if } p = \text{first}(p), \\ \text{offest}_2(p) & \text{if } p \neq \text{first}(p). \end{cases} \tag{218}$$

where

$$\text{offset}_1(p) \quad = \quad \sum_{\substack{p' < p: \\ p' = \text{first}(p')}} m_{k_{p'}}, \tag{219}$$

$$\text{offset}_2(p) \quad = \quad \sum_{\substack{p' \leq P: \\ p' = \text{first}(p')}} m_{k_{p'}} + \sum_{\substack{p' < p: \\ p' \neq \text{first}(p')}} m_{k_{p'}}. \tag{220}$$

The sorting matrix, $T_{\text{sort}}$, is an $M \times M$ matrix having elements

$$[T_{\text{sort}}]_{ij} = \begin{cases} 1 & \text{if } i = \text{sortedoffset}(p) + h \text{ and } j = \text{offset}(p) + h \text{ for some } p \in \mathcal{I}_P \text{ and } h \in \mathcal{I}_{m_{k_p}} - 1, \\ 0 & \text{otherwise,} \end{cases} \tag{221}$$

for $i, j \in \{0, \ldots, M - 1\}$. It is readily see that $T_{\text{sort}}$ is a permutation matrix. Consequently, the inverse of $T_{\text{sort}}$ is given by its transpose, i.e.,

$$T_{\text{sort}}^{-1} = T_{\text{sort}}^{\top}. \tag{222}$$

#### 8.5 Computation of the projection matrix $P$

Let $\Theta$ denote the gauge space spanned by the first $M-\gamma$ columns of $T^\dagger_{\text{dist}}$. It is readily seen that the resulting gauge-fixing projection matrix for $\Theta$ is given by

$$P = T^\dagger_{\text{dist}}\Big|_{M-\gamma} T^{-1\dagger}_{\text{dist}}, \tag{223}$$

where $|_{M-\gamma}$ denotes that the last $\gamma$ columns of a matrix have been set to zero. We note, however, that the gauge space $\Theta$ is not one of the parametric gauges discussed in our companion paper [1].

### 9 Observations motivating the conjecture

Based on the observations in the next two subsections, we conjecture that all allelic permutation models either (1) do not describe co-occurring alleles, (2) are equivalent to models that do not describe co-occurring alleles, or (3) have gauge freedoms.

#### 9.1 Single-orbit allelic models

**Claim 9.1.** *Single-orbit generalized one-hot models cannot describe co-occurring alleles.*

*Proof.* Every nontrivial orbit of a generalized one-hot model is defined by a set of generalized one-hot features,

$$\mathcal{O} = \left\{x^{c_1\ldots c_K}_{l_1\ldots l_K} : c_1,\ldots,c_K \in \mathcal{A}\right\}, \tag{224}$$

for some specified set of positions $\{l_1,\ldots,l_K\} \subseteq \mathcal{I}_L$, where each feature in the orbit $\mathcal{O}$ is given by

$$x^{c_1\ldots c_K}_{l_1\ldots l_K}(s) = \begin{cases} 1 & \text{if } c_k = s_{l_k} \text{ for } k=1,\ldots,K, \\ 0 & \text{otherwise,} \end{cases} \quad \text{for all} \quad s \in \mathcal{S}. \tag{225}$$

For any $s \in \mathcal{S}$, choosing $c_k = s_{l_k}$ for all $k = 1,\ldots,K$ yields a feature $x^{c_1\ldots c_K}_{l_1\ldots l_K} \in \mathcal{O}$ that is equal to 1 on $s$. Moreover, no other feature in $\mathcal{O}$ is nonzero when evaluated on $s$ because this would require that $s$ have a different character at one of the positions $l_k$. Therefore, for every sequence $s \in \mathcal{S}$, there is exactly one feature in the orbit $\mathcal{O}$ that is nonzero on $s$. This proves the claim. □

There are, however, are single-orbit allelic models that are not generalized one-hot models that describe co-occurring alleles. One example is the model based on the "two-hot" DNA embedding, in which each feature tests for the presence of one of two possible DNA bases:

$$\vec{x}^{\,\text{two}} = \begin{bmatrix} x^{\{\text{A,C}\}} \\ x^{\{\text{A,G}\}} \\ x^{\{\text{A,T}\}} \\ x^{\{\text{C,G}\}} \\ x^{\{\text{C,T}\}} \\ x^{\{\text{G,T}\}} \end{bmatrix} \quad \text{where} \quad x^{\{c_1,c_2\}}(c) = \begin{cases} 1 & \text{if } c \in \{c_1,c_2\}, \\ 0 & \text{otherwise} \end{cases} \quad \text{for all} \quad c \in \mathcal{A}_{\text{DNA}} \quad \text{and all} \quad \{c_1,c_2\} \subset \mathcal{A}_{\text{DNA}}. \tag{226}$$

This embedding comprises six allelic features. Since each DNA character is matched by three of the features, the embedding describes co-occurring alleles. This embedding is not equivalent to any generalized one-hot embedding, as it has the wrong dimension. In fact, the observation that $\vec{x}^{\,\text{two}}$ has six dimensions, transforms under a permutation representation, and is non-constant implies that it must transform under the representation

$$R^{\text{two}} \simeq 3R^{\text{triv}} \oplus R^{\text{sim}}. \tag{227}$$

The embedding $\vec{x}^{\,\text{two}}$ therefore has two gauge freedoms, which are readily seen to result from the three affine constraints $x^{\{\text{A,C}\}} + x^{\{\text{G,T}\}} = 1$, $x^{\{\text{A,G}\}} + x^{\{\text{C,T}\}} = 1$, and $x^{\{\text{A,T}\}} + x^{\{\text{C,G}\}} = 1$.

Another example of a single-orbit allelic embedding that describes co-occurring alleles is the "three-hot" DNA embedding, in which each feature tests for the presence of one of three possible DNA bases:

$$\vec{x}^{\,\text{three}} = \begin{bmatrix} x^{\{\text{A,C,G}\}} \\ x^{\{\text{A,C,T}\}} \\ x^{\{\text{A,G,T}\}} \\ x^{\{\text{C,G,T}\}} \end{bmatrix} \quad \text{where} \quad x^{\{c_1,c_2,c_3\}}(c) = \begin{cases} 1 & \text{if } c \in \{c_1,c_2,c_3\}, \\ 0 & \text{otherwise} \end{cases} \quad \text{for all} \quad c \in \mathcal{A}_{\text{DNA}} \quad \text{and all} \quad \{c_1,c_2,c_3\} \subset \mathcal{A}_{\text{DNA}}. \tag{228}$$

This embedding comprises four allelic features. Since each DNA character is matched by three of the features, the embedding describes co-occurring alleles. The fact that $\vec{x}^{\,\text{three}}$ has four dimensions, transforms under a permutation representation, and is non-constant implies that it must transform under the representation

$$R^{\text{two}} \simeq R^{\text{triv}} \oplus R^{\text{sim}} \simeq R^{\text{ohe}}. \tag{229}$$

The embedding $\vec{x}^{\,\text{three}}$ therefore has no gauge freedoms. Rather, it is equivalent to the one-hot embedding $\vec{x}^{\,\text{ohe}}$, i.e., equal up to a similarity transformation. Indeed, one can readily verify that

$$\vec{x}^{\,\text{three}} = A\vec{x}^{\,\text{ohe}}, \quad \text{where} \quad A = \begin{bmatrix} 0 & 1 & 1 & 1 \\ 1 & 0 & 1 & 1 \\ 1 & 1 & 0 & 1 \\ 1 & 1 & 1 & 0 \end{bmatrix}. \tag{230}$$

#### 9.2 Models defined by direct sums of direct products of single-position embeddings

Let us consider an embedding of the form

$$\vec{x}(s) = \bigoplus \vec{x}_{l_1 \dots l_k}, \quad \text{where} \quad \vec{x}_{l_1 \dots l_k} := \vec{x}_{l_1}(s) \otimes \dots \otimes \vec{x}_{l_k}(s), \tag{231}$$

the direct sum goes over an arbitrarily chosen set of position sets $\{l_1, \dots, l_k\}$, and each $\vec{x}_{l_j} = (x_{l_1}^{i_1}, \dots, x_{l_k}^{i_k})$ is an arbitrary single-position sequence embedding, where each $x_{l_j}^{i_j}$ is a function that only depends on sequence $s$ at position $l_j$. We assume that none of the single-position features are constant over all characters, i.e. if $x_l^i$ denotes the $i$th feature in the single-position embedding $\vec{x}_l$ then $x_l^i(c)$, $c \in \mathcal{A}$, is not a constant function. We note that the constant feature can be part of embedding $\vec{x}$, it corresponds to the empty set in the direct sum over position sets in (231).

Let $\gamma$ denote the number of gauge freedoms of the model that uses embedding $\vec{x}$, and let $\gamma_{\text{goh}}$ denote the number of gauge freedoms of the generalized one-hot model that uses the embedding of the same direct sum/direct product structure as $\vec{x}$, given in (231), but with $\vec{x}_{l_i} = \vec{x}_{l_i}^{\,\text{ohe}}$ for all $l_i$.

**Claim 9.2.** *If a model built from embedding $\vec{x}$ given in formula (231) transforms under a permutation representation of $H_{\text{PSCP}}$, then the model has at least as many gauge freedoms as the corresponding one-hot model, i.e.*

$$\gamma \geq \gamma_{\text{goh}}. \tag{232}$$

*Proof.* Let us assume we have an embedding $\vec{x}$ that satisfies the assumptions above, and let $R$ be the permutation representation under which it transforms. For each $h$, $R(h)$ is a permutation matrix, and therefore induces a permutation on the set of features which we denote by $h'$. With this notation, the equivariance-defining equation becomes

$$\vec{x}(h(s)) = h'(\vec{x}(s)), \quad s \in \mathcal{S}, h \in H_{\text{PSCP}}. \tag{233}$$

We first show that, for each $h \in H_{\text{PSCP}}$, $h'$ does not mix features corresponding to different position sets, i.e. $h'$ maps each feature vector $\vec{x}_{l_1 \dots l_k}$ onto itself. First, if the embedding contains the constant feature $x_0$, we have $h'(x_0(s)) = x_0(h(s))$ =constant for all $s$, therefore $x_0$ is mapped to itself. (We note that we assumed that there is at most one constant feature in the embedding $\vec{x}$.). Next, let us consider how $h'$ acts on single-position features. Let $x_l^i$ denote the $i$th coordinate of the singe-position embedding $\vec{x}_l$. If $x_l^i$ is mapped to a possibly higher order feature $x_{l_1 \dots l_k}^{i_1 \dots i_k}$ ($k \geq 1$), then using the product form $h = h_1 \cdots h_L$, we have

$$x_l^i(h(s)) = x_{l_1 \dots l_k}^{i_1 \dots i_k}(s) = x_{l_1}^{i_1}(s) \cdot \ldots \cdot x_{l_k}^{i_k}(s). \tag{234}$$

We see that the left-hand side function only depends on a sequence's character at position $l$. Therefore, for any position $l_j \neq l$, if we change $s$ at $l_j$, the left-hand side of the equation above does not change, while the right hand side does. Note that here we use the assumption that none of the single-position features $x_{l_j}^{i_j}(s)$ are constant functions of $s$. Therefore, we see that, to avoid contradiction, we need to have $k = 1$ and $l_1 = l$, that is, any single-position feature vector $\vec{x}_l(s)$ gets mapped onto itself by $h'$. Finally, consider how $h'$ acts on higher order features. By the definition of higher order feature vectors as tensor products of single-position feature vectors, and using the product form $h = h_1 \cdots h_L$, we obtain

$$x_{l_1 \dots l_k}^{i_1 \dots i_k}(h(s)) = \prod_{j=1}^{k} x_{l_j}^{i_j}(h(s)) = \prod_{j=1}^{k} x_{l_j}^{i_j}(h_{l_j}(s)) = \prod_{j=1}^{k} x_{l_j}^{i'_j}(s) = x_{l_1 \dots l_k}^{i'_1 \dots i'_k}(s), \tag{235}$$

where $x_{l_j}^{i'_{l_j}} := h'_{l_j}(x_{l_j}^{i_{l_j}})$. We thus find that $\vec{x}_{l_1 \dots l_k}$ also gets mapped onto itself by $h'$.

We conclude that for each $h \in H$, the permutation matrix $R(h)$ is block-diagonal with blocks $R_{l_1 \ldots l_k}(h)$ corresponding to the position sets $l_1 < \ldots < l_k$ in formula (231). Therefore

$$\vec{x}(h(s)) = \bigoplus_{l_1 < \ldots < l_k} \vec{x}_{l_1 \ldots l_k}(h(s)) = \bigoplus_{l_1 < \ldots < l_k} R_{l_1 \ldots l_k}(h) \vec{x}_{l_1 \ldots l_k}(s). \tag{236}$$

Considering a term in the right hand side expression above, we saw in equation (235) that how $R_{l_1 \ldots l_k}(h)$ acts on $\vec{x}_{l_1 \ldots l_k}$ is completely determined by how it acts on single-site feature vectors, specifically, $R_{l_1 \ldots l_k}(h) = R_{l_1}(h) \otimes \ldots \otimes R_{l_k}(h)$, where $R_{l_j}(h)$ is the block of $R(h)$ that acts on the single-position feature vector $\vec{x}_{l_j}$. In summary, we found that

$$R = \bigoplus R_{l_1} \otimes \ldots \otimes R_{l_k}, \tag{237}$$

where each $R_{l_i}$ is permutation representation of $S_\alpha$.

Let us look at the Maschke decomposition of a factor $R_{l_j}$ in the expression above. A permutation representation is reducible, in particular, its Maschke decomposition always contains the trivial representation. From Claim 4.2, we further know that the Maschke decomposition of $R_{l_j}$ can only contain copies of the trivial representation $R^{\text{triv}}$ and copies of the simplex representation $R^{\text{sim}}$ with degree $\alpha - 1$. But if it were to contain copies of only the trivial representation, then $\vec{x}_{l_j}$ would be the trivial embedding which would contradict our assumption that none of the first order features are constant over all characters. This means that the Maschke decomposition of $R_{l_j}$ also has to contain a copy of the simplex representation. Therefore the Maschke decomposition of $R_{l_j}$ looks like

$$R_{l_j} \simeq Q^{\text{triv}} R^{\text{triv}} \oplus Q^{\text{sim}} R^{\text{sim}}, \tag{238}$$

where $Q^{\text{triv}} \geq 1$ and $Q^{\text{sim}} \geq 1$.

Since $Q^{\text{triv}} = 1$ and $Q^{\text{sim}} = 1$ in the Maschke decomposition of the corresponding one-hot model, we see that whenever a coordinate of $\vec{x}$ is zeroed out in the embedding distillation procedure, so is the corresponding coordinate of $\vec{x}^{\text{ohe}}$, and therefore, $\gamma \geq \gamma_{\text{goh}}$. □
